# The methyltransferases METTL7A and METTL7B confer resistance to thiol-based histone deacetylase inhibitors

**DOI:** 10.1101/2022.10.07.511310

**Authors:** Robert W. Robey, Christina M. Fitzsimmons, Wilfried M. Guiblet, William J.E. Frye, José M. González Dalmasy, Li Wang, Drake A. Russell, Lyn M. Huff, Andrew J. Perciaccante, Fatima Ali-Rahmani, Crystal C. Lipsey, Heidi M. Wade, Allison V. Mitchell, Siddhardha S. Maligireddy, David Terrero, Donna Butcher, Elijah F. Edmondson, Lisa M. Jenkins, Tatiana Nikitina, Victor B. Zhurkin, Amit K. Tiwari, Anthony D. Piscopio, Rheem A. Totah, Susan E. Bates, H. Efsun Arda, Michael M. Gottesman, Pedro J. Batista

## Abstract

Histone deacetylase inhibitors (HDACis) are part of a growing class of epigenetic therapies used for the treatment of cancer. While elevated levels of the efflux pump P-gp are associated with *in vitro* resistance to romidepsin, this mechanism does not translate to the clinic. We developed a romidepsin-resistant cell line with a resistance mechanism independent of P-gp function that acts upstream of the deacetylation process. We found that expression of the methyltransferase METTL7A is necessary for resistance, and that expression of METTL7A in naïve cells can drive resistance to thiol-containing HDACis. We demonstrate that METTL7A can methylate romidesin *in vitro* and that the ability of METTL7A to drive resistance to thiol-containing HDACis can be blocked by the methyltransferase inhibitor DCMB. Our data supports a model whereby exposure of cells to romidepsin selects for upregulation of the methyltransferase METTL7A, which in turn modifies the zinc-binding thiol, inactivating the drug.

## INTRODUCTION

Post-translational modification of histone tails via methylation, phosphorylation, or acetylation is an essential aspect of gene regulation in the cell [1]. Histone acetylation is controlled by the interplay between histone acetyltransferases (HATs) that catalyze the addition of acetyl groups to histones, and histone deacetylases (HDACs) that catalyze their removal. Increased histone acetylation leads to a more accessible chromatin configuration and increased gene transcription, while decreased histone acetylation is associated with transcriptional repression [2]. Alterations in the expression of HDAC proteins have been demonstrated in various cancers, potentially leading to altered transcriptional regulation of genes related to cell growth and differentiation [3]. Several HDAC inhibitors (HDACis) have been developed that can block histone deacetylation, thus resulting in increased histone acetylation via unrestrained HATs activity [4].

In clinical trials, HDACis have shown immense promise in the treatment of T-cell lymphoma. Two HDACis, vorinostat and romidepsin, have been approved as single agents by the United States Food and Drug Administration (FDA) for the treatment of cutaneous T-cell lymphoma [5–8]. Additionally, belinostat was approved as a single agent for the treatment of peripheral T-cell lymphoma [9] and tucidinostat (chidamide) has been approved in China as a single agent for the treatment of peripheral T-cell lymphoma [10]. Romidepsin is unique among the HDACis that have been approved for cancer treatment in that it is a bicyclic depsipeptide with an internal disulfide bond that must be reduced to form the activated molecule [11]. The reduced form of romidepsin has two free sulfhydryl groups, one of which can coordinate with zinc in the active site of HDACis, thus reversibly inhibiting the enzymes [11].

Despite dramatic responses to treatment in some T-cell lymphoma patients, romidepsin can lose effectiveness after the initial response period. In addition, clinical trials with solid tumors have been unsuccessful, suggesting intrinsic mechanisms of resistance. Romidepsin was identified as a substrate of the ATP binding cassette transporter P-glycoprotein (P-gp, encoded by the *ABCB1* gene) based on early studies with the NCI-60 cell lines. Several groups have reported overexpression of P-gp in cell lines that are selected for resistance to romidepsin [12–16]. However, in clinical samples, expression levels of *ABCB1* did not predict response to romidepsin [17], prompting us to search for non-P-gp mechanisms of resistance.

To generate a solid tumor model of romidepsin resistance, MCF-7 breast cancer cells were selected with romidepsin and verapamil, an inhibitor of P-gp, to prevent the emergence of P-gp as a resistance mechanism. While the resulting cell line, MCF-7 DpVp300, was selectively resistant to romidepsin, it did not overexpress P-gp. As such, resistance of DpVp300 cells to romidepsin is not driven by P-gp. We performed Assay for Transposase-Accessible Chromatin sequencing (ATAC-seq) and RNA-sequencing (RNA-seq) on MCF-7 and MCF-7 DpVp300 cells to identify possible mechanisms of resistance. One of the genes that was most upregulated in MCF-7 DpVp300 cells compared to MCF-7 cells was *METTL7A*, a putative methyltransferase. This gene was also found to be upregulated in HuT78 DpVp50 and HuT78 DpP75 cells, two romidepsin-resistant T-cell lymphoma cell lines we had generated previously [18]. Here, we demonstrate that METTL7A overexpression can confer resistance not only to romidepsin, but to other HDACis that have a thiol as the zinc-binding moiety. Expression of the close homolog METTL7B can also confer resistance to HDACis bearing a thiol group, albeit with less efficiency. Overexpression of these methyltransferases represents a previously unknown mechanism of resistance to thiol-based HDACis and a potential target to improve HDACi-based therapies.

## RESULTS

### MCF-7 DpVp300 cells are selectively resistant to romidepsin

The MCF-7 DpVp300 cell line was generated via prolonged selection of MCF-7 cells with romidepsin in the presence of the P-gp inhibitor verapamil to prevent any selection benefit from P-gp overexpression. To determine which drugs the MCF-7 DpVp300 cells are sensitive or resistant to (the cross-resistance profile), cytotoxicity assays were performed with several chemotherapeutic agents, including HDACis. Interestingly, the DpVp300 cell line was much more resistant to romidepsin (Fig. 1A, on average 218-fold more resistant) and much less resistant to other HDACis, such as belinostat (8-fold), panobinostat (2.5-fold), and vorinostat (5-fold). Additionally, the MCF-7 DpVp300 line was not cross resistant to other chemotherapeutic agents such as 5-FU, methotrexate, teniposide, or topotecan, exhibiting no more than 3-fold more resistance to those compounds than MCF-7 cells (Supplemental Table S1).

**Fig. 1.**
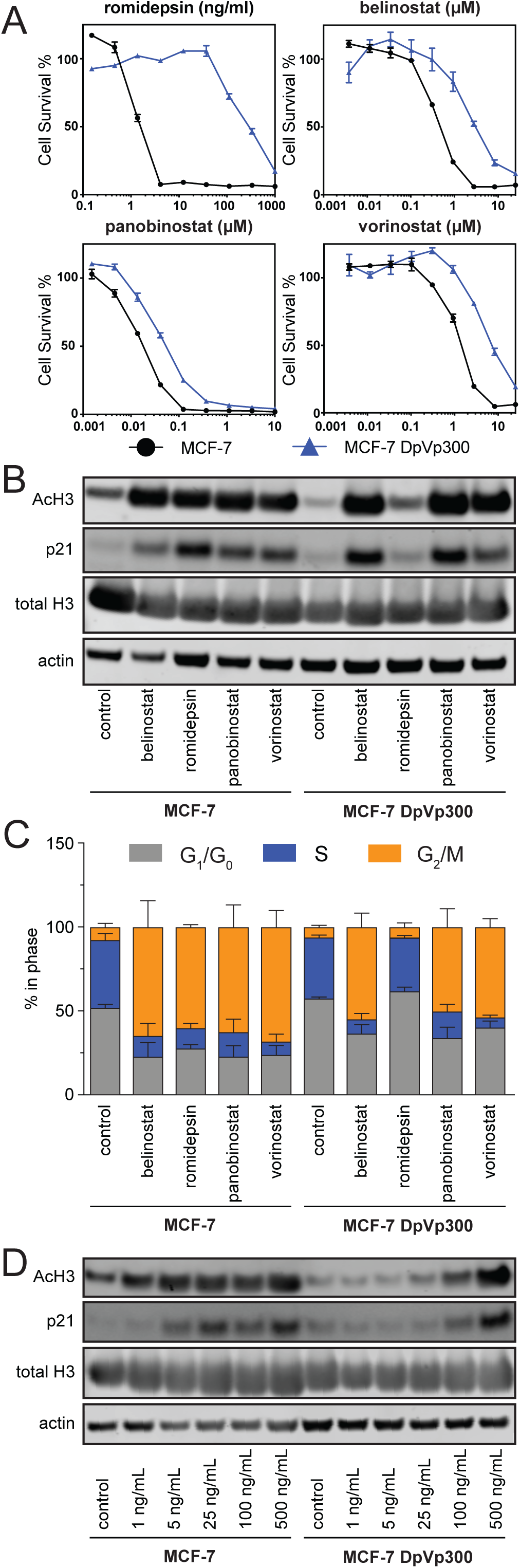
MCF-7 DpVp300 cells are selectively resistant to romidepsin. (A) Three-day cytotoxicity assays were performed on MCF-7 cells (black circles) and MCF-7 DpVp300 cells (blue triangles) with the histone deacetylase inhibitors romidepsin, belinostat, panibinostat and vorinostat. (B) MCF-7 and MCF-7 DpVp300 cells were plated and allowed to attach overnight. Cells were subsequently treated with belinostat (5 µM), romidepsin (10 ng/ml), panobinostat (5 µM), or vorinostat (5 µM) for 24 h after which protein was extracted and subjected to immunoblot analysis. Membranes were incubated with antibodies to pan acetylated histone H3 (AcH3), total histone H3 (total H3), p21 or actin. One of three independent experiments is shown. (C) MCF-7 and MCF-7 DpVp300 cells were plated and allowed to attach overnight after which cells were exposed to belinostat (5 µM), romidepsin (10 ng/ml), panobinostat (5 µM), or vorinostat (5 µM) for 24 h. Cells were subsequently analyzed by flow cytometry and the percentage of cells in each phase of the cell cycle was determined using ModFit LT v 5.0. The graph was generated from data compiled from three independent experiments. Error bars are standard deviation. (D). MCF-7 and MCF-7 DpVp300 cells were plated and allowed to attach overnight after which cells were exposed to increasing concentrations of romidepsin for 24 h. After harvesting cells and subjecting extracted protein to immunoblot analysis, proteins were probed with antibodies as described in (B). One of three independent experiments is shown.

As HDACi treatment results in increased levels of acetylated histones and p21 transcription [19], we examined the effect of HDACis on global histone acetylation and p21 levels. MCF-7 and MCF-7 DpVp300 cells were treated with romidepsin, belinostat, panobinostat, or vorinostat for 24 h after which protein was extracted and subjected to analysis by immunoblot. MCF-7 cells treated with any of the HDACis displayed increased levels of acetylated histone H3 and p21 compared to untreated cells (Fig. 1B). In contrast, MCF-7 DpVp300 cells treated with romidepsin did not exhibit the increased levels of acetylated histone H3 or p21 that were found with MCF-7 cells; however, treatment with the other HDACis resulted in higher levels of acetylated histone H3, which suggested a resistance mechanism that is specific to romidepsin and acts upstream of the deacetylation process.

Treatment of cancer cell lines with HDACis is known to cause cell cycle perturbations, with some cell lines demonstrating a G_1_/G_0_ arrest and others a G_2_/M arrest, depending upon which checkpoints were most intact [20–22]. Cell cycle analysis was performed on MCF-7 and DpVp300 cells treated with belinostat, romidepsin, panobinostat, or vorinostat for 24h as outlined above. In the MCF-7 cells treated with the HDACis, a decrease of cells in the G_1_/G_0_ phase and an increase in cells in the G_2_/M phase was observed (Supplemental Fig. S1A). While a similar pattern was observed in MCF-7 DpVp300 cells when treated with belinostat, panobinostat or vorinostat, no such changes were observed with romidepsin treatment. When data from three independent experiments were compiled (Fig. 1C), we observed a significant decrease in the percentage of cells in the G_1_/G_0_ and S phases of the cell cycle and an increase of cells in the G_2_/M phase in MCF-7 cells treated with any of the HDACis. Similar changes were observed when DpVp300 cells were treated with belinostat, panobinostat or vorinostat; however, romidepsin treatment did not produce significant changes (Fig. 1C).

To examine the effects of higher doses of romidepsin, we treated MCF-7 and DpVp300 cells with increasing concentrations of romidepsin for 24 h and examined levels of histone acetylation and p21 by immunoblot. We observed that MCF-7 cells displayed increased levels of histone H3 acetylation when treated with as little as 1 ng/ml romidepsin (Fig. 1D). In contrast, increased histone acetylation was not observed until MCF-7 DpVp300 cells were treated with at least 100 ng/ml romidepsin. Based on these results, we concluded the resistance mechanism present in MCF-7 DpVp300 cells protected the cells from romidepsin levels at least 100 times higher than MCF-7 cells – consistent with the earlier observation in cytotoxicity assays that MCF-7 DpVp300 cells are at least 200-fold resistant to romidepsin (Fig. 1A).

### P-gp is not responsible for romidepsin resistance in MCF-7 DpVp300 cells

We and others have demonstrated that selection with romidepsin as a single agent leads to overexpression of P-gp as a resistance mechanism [13, 15, 16, 23]. The MCF-7 DpVp300 line was generated to identify alternative mechanisms of resistance, blocking P-gp activity with verapamil to prevent its emergence as a resistance mechanism. To confirm that P-gp does not play a role in romidepsin resistance in the DpVp300 cell line, surface expression of P-gp was measured by flow cytometry using the UIC2 antibody. P-gp was readily detected in the *ABCB1*-transfected MDR-19 cell line, as shown by the difference in fluorescence between the isotype control peak (solid line) and UIC-2 peak (dashed line) (Supplemental Fig. S1B). However, P-gp was largely undetected in the empty vector transfected line, as shown by the overlapping isotype control (solid line) and UIC-2 peak (dashed line). We did not detect P-gp expression in either the MCF-7 cells or in MCF-7 DpVp300 cells, as shown by the overlapping isotype control (solid line) and UIC-2 (dashed line) peaks. Cross-resistance to romidepsin in the MDR-19 line compared to empty vector transfected cells leads to approximately the same degree of resistance as that observed between DpVp300 cells and MCF-7 cells (Supplemental Fig. S1C). Thus, we concluded the mechanism of resistance at work in the MCF-7 DpVp300 cells is P-gp-independent.

### MCF-7 DpVp300 cells have significant changes in gene expression

Both MCF-7 DpVp300 and HuT78 DpVp50 cells, which we previously generated [18], have acquired mechanisms of resistance to romidepsin that do not involve P-gp/ABCB1 function. To determine which molecular pathways might be involved in the resistance mechanism we assessed changes in gene expression between the sensitive and the drug-resistant cells.

We observed significant changes in steady-state mRNA levels between the sensitive and DpVp cell lines (Fig. 2A) for both MCF-7 and Hut78. Approximately 500 genes were upregulated, and 180 genes were downregulated (|l2fc| > 1.0 and padj < 0.05) in both HuT78 and MCF-7 DpVp cells. The majority of GO-terms enriched for up-or down-regulated genes were distinct between the two cell lines and offered no indication of underlying genes or molecular pathways that could drive resistance to romidepsin (Supplemental Fig. S2A). As expected, we observed little change in the expression level of *ABCB1* mRNA. We focused our analysis on the genes most up-regulated in both the MCF-7 and HutT78 cells. We identified 5 genes that are highly up-regulated in both DpVp variants (log2 fold change >= 5.0 and padj < 1e-300) (Fig. 2B). These genes include *PEG10*, *PRAME*, *SEMA3A*, *SGCE*, and *METTL7A*. We decided to focus on METTL7A, a putative methyltransferase that has not been extensively studied but has been linked to lipid droplet formation in cells [24]. METTL7B, a paralog of METTL7A, is an alkyl-thiol methyltransferase that is capable of methylating thiol groups of various compounds such as captopril, hydrogen sulfide, and dithiothreitol [25]. As the active form of romidepsin contains a thiol as the zinc binding group, and methylation of the thiol group would prevent coordination of the molecule with zinc in the HDAC binding pocket, we hypothesized that METTL7A might be able to inactivate romidepsin or other HDACis with a thiol as the zinc-binding group. Immunoblot analysis confirmed high levels of METTL7A protein not only in MCF-7 DpVp300 and HUT78 DpVp50 cells but also in another resistant cell line, HuT78 DpP75 [18] (Fig. 2C). Taken together, these results indicate that METTL7A is upregulated in both MCF-7 DpVp300 and HuT78 DpVp50 and may play a role in romidepsin resistance.

**Fig. 2.**
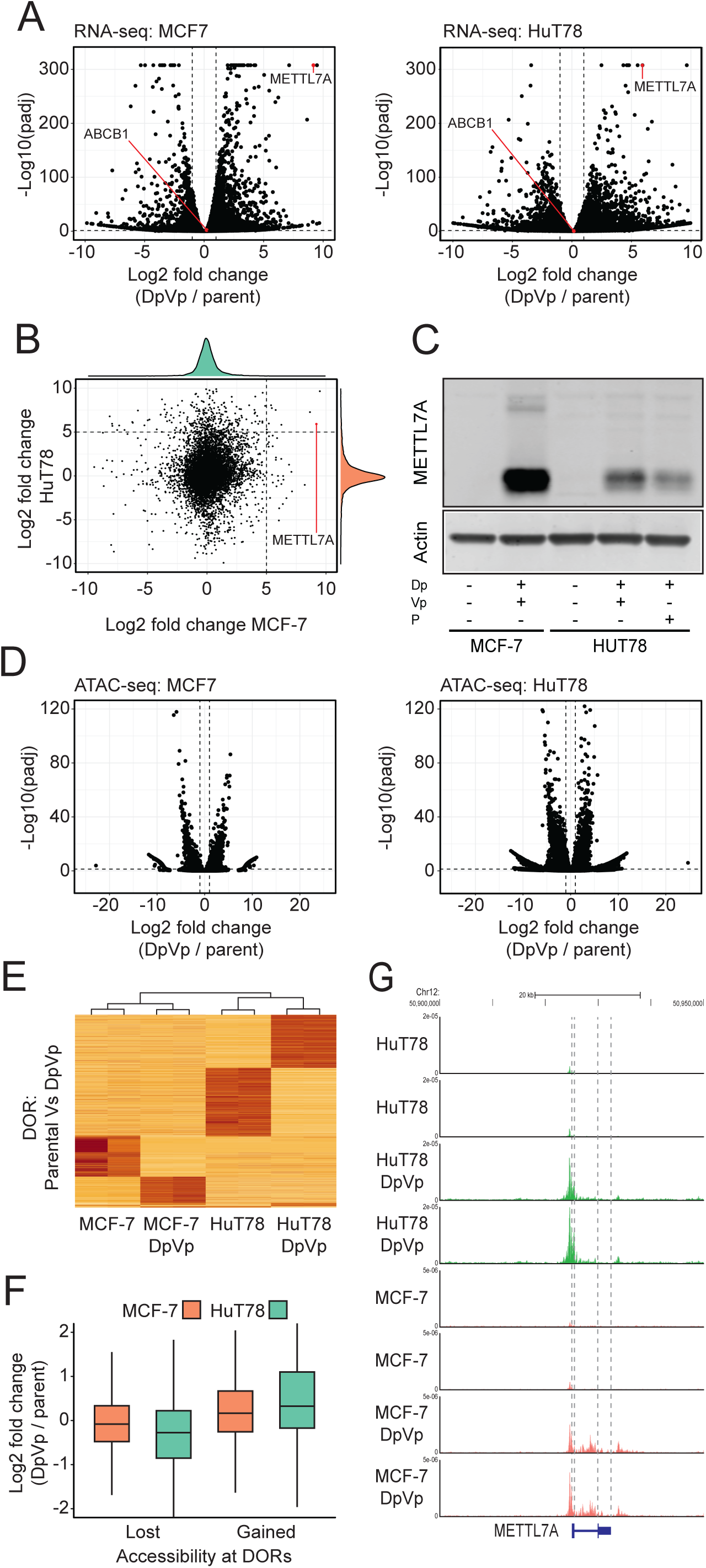
RNA Seq and ATAC Seq analysis of MCF-7 and MCF-7 DpVp300 cells. (A) Volcano plot depicting steady state fold change in mRNA levels between MCF-7 and MCF-7 DpVp300 (left) and HuT78 and HuT78 DpVp50 (right). Changes in gene expression were determined with DEseq2. The drug efflux gene *ABCB1* and *METTL7A* are indicated with a red line Scatterplot depicting the log2 fold change of transcripts in MCF-7 and MCF-7 DpVp300 compared to the log2 fold change in HuT78 and HuT78 DpVp50 cells. A log2 fold change of 5 is indicated on the x and y-axis with a dashed line. *METTL7A* is indicated with a red line. (B) Immunoblot of METTL7A in the MCF-7 and HuT78 resistance models. (C) Volcano plot depicting the fold change of differentially open regions in MCF7 cells (left) and Hut78 cells (right) identified with PEPATAC. (D) Heatmap depicting changes in the differentially open regions in MCF-7, MCF-7 DpVp300, HuT78 and HuT78 DpVp50 cells. (E) Box plot showing log2 fold change in mRNA levels for genes linked to differentially open region that gain or lose accessibility in DpVp cells. Genes linked to differentially open regions with both gain and loss of accessibility are excluded. (F) Differentially open regions at the genomic locus of METTL7A in MCF-7, MCF-7 DpVp300, HuT78, and HuT78 DpVp50 cells.

### METTL7A is upregulated at the transcriptional level

Several mechanisms could drive upregulation of METTL7A in DpVp cells. To determine if expression of METTL7A is regulated at the level of transcription, we compared chromatin accessibility in the parent and the DpVp lines for MCF-7 and HuT78. A total of 19,937 differentially open regions (DORs) (padj < 0.05) in the MCF-7 cells and 34,737 DORs in the HuT78 cells were identified (Fig. 2D). Consistent with our observation that there is little overlap between up- and down-regulated genes at the mRNA level, we also observe little overlap between MCF-7 and HuT78 DpVp DORs, with only 737 DORs common to both DpVp cell lines (Fig. 2E). *De novo* motif discovery of transcriptional binding sites at DORs shows little overlap between the two cell lines, (Supplemental Fig. S2B), suggesting different transcriptional networks are active in the two DpVp cell lines. As expected, genes associated with highly accessible DORs tend to be upregulated in the DpVp cells (Fig. 2F). Finally, upregulation of METTL7A is associated with open DORs at the genomic locus (Figure 2G) and an increase in density of subnucleosomal DNA fragments (Supplemental Fig 2C). The mRNA coding for METTL7A has been shown to undergo extensive editing at the 3’ end, and protein expression has been shown to be suppressed by adenosine deaminase acting on RNA (ADAR1/2) and miR-27a [32]. We observe extensive editing of the 3’-UTR of METTL7A in both of our resistance models (Supplemental Fig. S2D), suggesting that post-transcriptional regulation might also play a role in the upregulation of METTL7A.

### MCF-7 DpVp300 cells are resistant to other thiol-based HDACIs

Since METTL7B has been shown to be an alkyl-thiol methyltransferase, we hypothesized that if METTL7A is the driver of romidepsin resistance, the MCF-7 DpVp300 might be resistant to other HDACis with a thiol group. We performed cytotoxicity assays on MCF-7 and MCF-7 DpVp300 cells with largazole, a natural-product HDACi [26]; OKI-005, an HDACi based on the structure of largazole and currently in clinical trials [27]; KD5170, a class I and II HDACi [28]; and NCH-51, an HDACi based on the structure of vorinostat [29]. All of these small molecules are thioester prodrugs that hydrolyze to yield a thiol as the zinc binding group. We found that the DpVp300 cells were resistant to all the thiol-based HDACis tested (Fig. 3A and Supplementary Table S2).

**Fig. 3.**
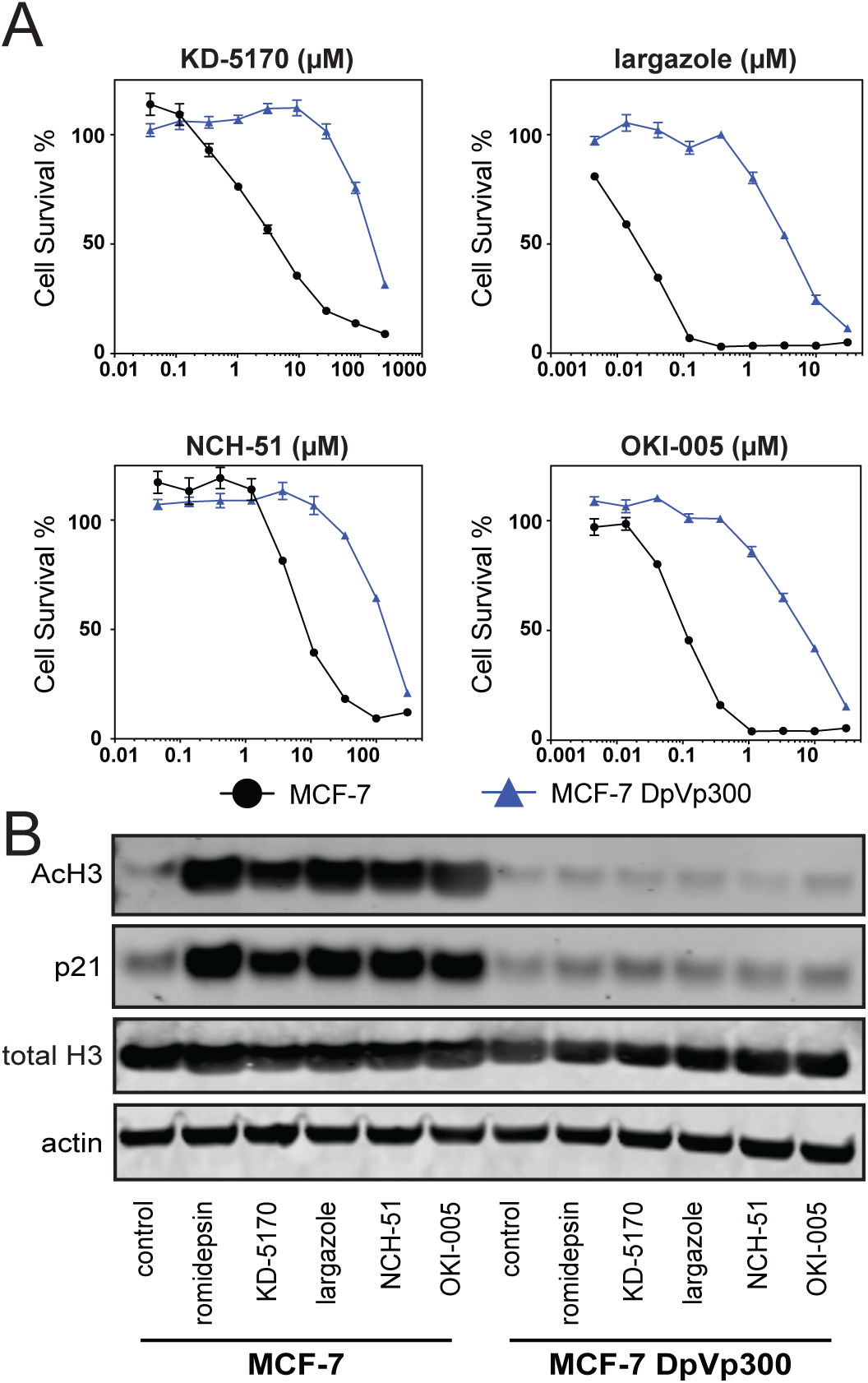
METTL7A overexpression confers resistance to thiol-based HDACIs in the MCF-7 DpVp300 line. (A) Three-day cytotoxicity assays were performed on MCF-7 cells (black circles) and MCF-7 DpVp300 cells (blue triangles) with KD-5170, largazole, NCH-51, and OKI-005. (B) MCF-7 and MCF-7 DpVp300 cells were plated and allowed to attach overnight. Cells were subsequently treated with romidepsin (10 ng/ml), KD-5170 (10 µM), largazole (250 nM), NCH-51 (25 µM), or OKI-005 (500 nM) for 24 h after which protein was extracted and subjected to immunoblot. Blots were incubated with antibodies to pan acetylated histone H3 (AcH3), total histone H3 (total H3), p21 or actin. Results from one of three independent experiments are shown.

Immunoblot analysis of histone acetylation and p21 expression showed that while acetylated histone H3 and p21 expression increased when MCF-7 cells were treated with any of the above thiol-based HDACis, no increase in expression of these proteins was observed in DpVp300 cells treated with any of the HDACis (Fig. 3B). This suggests that MCF-7 DpVp300 cells were selectively resistant to the effects of thiol-based HDACis. Further, these results support the hypothesis that METTL7A methylates and inactivates thiol-based HDACis, driving resistance to treatment.

### Knockout of METTL7A in MCF-7 DpVp300 cells restores sensitivity to thiol-based HDACIs

To investigate whether METTL7A was responsible for the observed resistance to romidepsin and other HDACis with an active thiol group, we performed CRISPR-Cas9-mediated knockout of *METTL7A*. We generated two clones that were completely negative for METTL7A expression, 7A KO B1 and 7A KO B3, as well as one clone that had reduced expression but not total loss of METTL7 protein, 7A KO B4 (Fig. 4A). When the clones were treated with romidepsin for 24 h, the ability of romidepsin to induce histone H3 acetylation and p21 was restored in the complete knockouts while little effect was observed in the clone with partial knockdown (Fig. 4A). In cytotoxicity assays, sensitivity to all thiol-based HDACis was restored nearly to the level of parent MCF-7 cells in the knockout clones, while resistance to the HDACis was only slightly decreased in the partial knockdown clone (Fig. 4B). Sensitivity to vorinostat, which does not contain a thiol group, was unchanged (Supplemental Table S3).

**Fig. 4.**
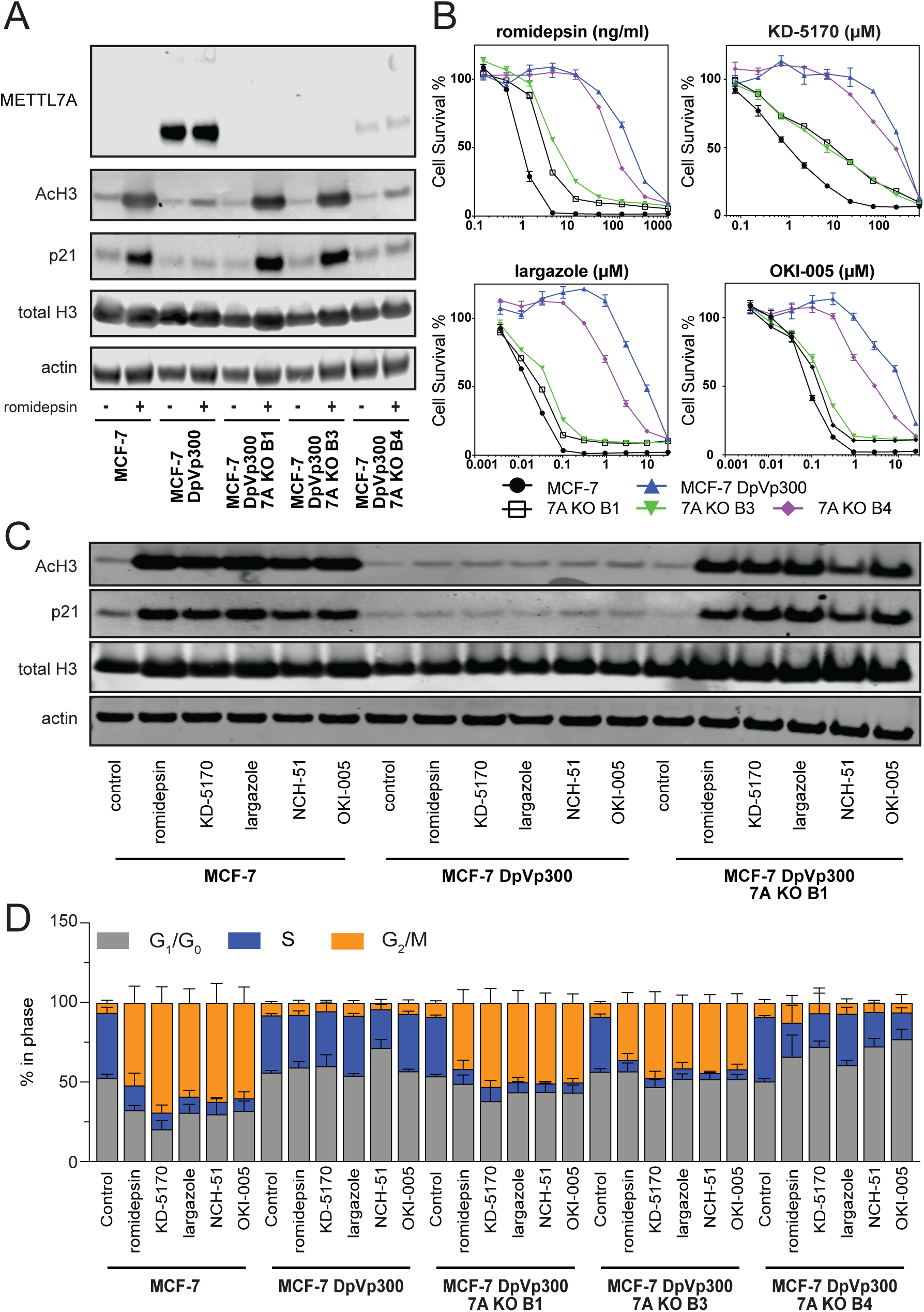
Deletion of METTL7A restores sensitivity to thiol-based HDACIs in MCF-7 DpVp300 cells. (A) MCF-7 and MCF-7 DpVp300 cells, as well as knockout clones (7A KO B1 and 7A KO B3) and partial knockdown clone 7A KO B4 were plated and allowed to attach overnight. Cells were subsequently left untreated or treated with romidepsin (10 ng/ml) for 24 h. After harvesting cells and subjecting extracted protein to PAGE, proteins were transferred to nitrocellulose that was probed with METTL7A antibody as well as pan acetylated histone H3 (AcH3), total histone H3 (total H3), p21 or actin antibodies. (B) Three-day cytotoxicity assays were performed with romidepsin, KD-5170, largazole, NCH-51, OKI-005 and vorinostat on MCF-7 cells (black circles), MCF-7 DpVp300 cells (blue triangles), 7A KO B1 cells (open black squares), 7A KO B3 (inverted green triangles) and 7A KO B4 (purple diamonds). (C) MCF-7, MCF-7 DpVp300 and 7A KO B1 cells were plated and allowed to attach overnight. Cells were subsequently treated with romidepsin (10 ng/ml), KD-5170 (10 µM), largazole (250 nM), NCH-51 (25 µM), or OKI-005 (500 nM) for 24 h after which protein was extracted and subjected to PAGE. Proteins were transferred to nitrocellulose membranes and blots were incubated with antibodies to pan acetylated histone H3 (AcH3), total histone H3 (total H3), p21 or actin. Results from one of three independent experiments is shown. (D) MCF-7, MCF-7 DpVp300 and 7A KO B1 cells were plated and allowed to attach overnight after which cells were exposed to belinostat (5 µM), romidepsin (10 ng/ml), panobinostat (5 µM), or vorinostat (5 µM) for 24 h. Cell cycle analysis was subsequently performed and the percentage of cells in each phase was determined using ModFit LT v 5.0. The graph was generated from data compiled from three independent experiments. Error bars indicate standard deviation.

MCF-7, MCF-7 DpVp300 and 7A KO B1 cells were subsequently treated with romidepsin, KD-5170, largazole, NCH-51, or OKI-005 for 24 h, after which the effects on histone acetylation and p21 expression were examined by immunoblot (Fig. 4C). While MCF-7 cells displayed increases in histone acetylation and p21 after treatment with any of the HDACis, no increases were observed in MCF-7 DpVp300 cells, and knockout of METTL7A restored the ability of the thiol-based HDACis to induce histone acetylation and p21.

In addition, we performed cell cycle analysis on MCF-7, MCF-7 DpVp300 cells and the 3 knockout clones (Fig. 4D and Supplemental Fig. S3). Cells were treated at the concentrations used above for the immunoblot for 24 hrs. We observed that MCF-7 cells had fewer cells in the G_1_/G_0_ and S phases of the cell cycle and an increased number of cells in G_2_/M phase after treatment with any of the compounds (Fig. 4D). Changes in the cell cycle were not observed in DpVp300 cells treated with any of the drugs except in the case of NCH-51. In agreement with the cytotoxicity data, we observed significant cell cycle perturbations after treatment with the thiol-based HDACis in both METTL7A knockout clones. These effects were comparable to what we observed with MCF-7 cells. In the partial knockout clone, we noted some significant cell-cycle changes; however, these changes were less pronounced compared to MCF-7 cells, suggesting that even a low level of METTL7 expression is still protective to the cells. These results all pointed to METTL7A as a mediator of resistance to HDACis with a thiol as the zinc-binding moiety.

### Overexpression of METTL7A or METTL7B confers resistance to thiol-based HDACis

To further test the ability of METTL7A to confer resistance to thiol-containing HDACis, we transfected HEK-293 cells, which are sensitive to HDACis, with *METTL7A* and the close paralog *METTL7B*. We selected two clones of each with comparable protein levels of METTL7A or METTL7B (Fig. 5A) and performed three-day cytotoxicity assays with largazole, OKI-005, romidepsin, NCH-51 or KD5170 along with two non-thiol HDAC inhibitors, vorinostat and mocetinostat. METTL7A overexpression conferred resistance to all thiol-based HDACis tested (Fig. 5B and Supplemental Table S4) compared to empty vector transfected cells (Vector). METTL7B overexpression conferred low-level resistance to romidepsin and it also appeared to be less effective than METTL7A for largazole and KD5170 (Fig. 5B).

**Fig. 5.**
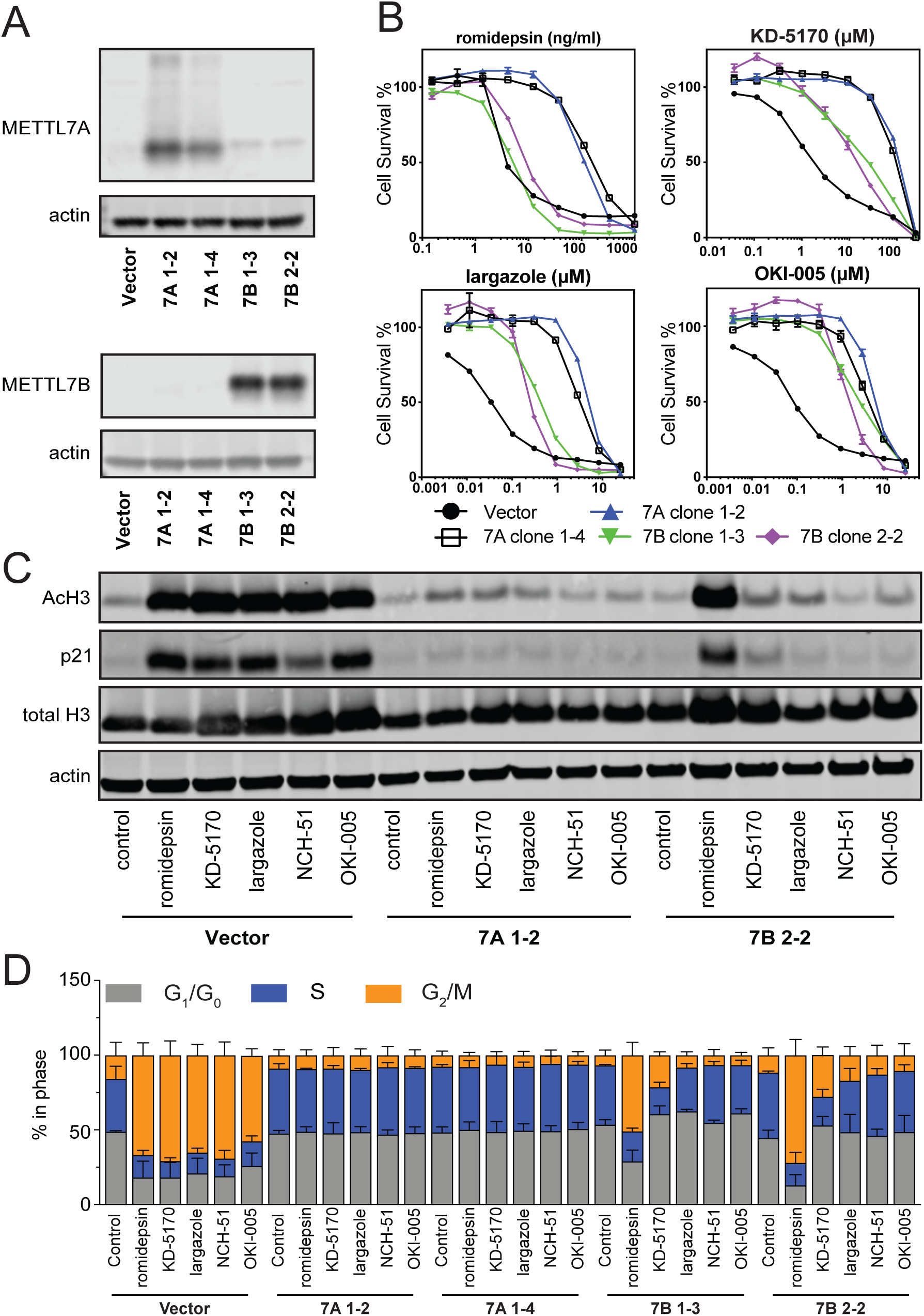
METTL7A and METTL7B overexpression confers varying levels of resistance to thiol-based HDACis. (A) HEK293 cells were transfected with empty vector (vector) or with vector encoding METTL7A (clones 7A 1-2 and 7A 1-4) or METTL7B (clones 7B 1-3 and 7B 2-2). Proteins were extracted and subjected to immunoblot. The blots were probed with antibodies to METTL7A, METTL7B or actin. (B) Three-day cytoptoxicity assays were performed with romidepsin, KD-5170, largazole, or OKI-005 on empty vector (vector) cells (black circles), 7A 1-2 cells (blue triangles), 7A 1-4 cells (open black squares), 7B 1-3 (inverted green triangles) and 7B 2-2 (purple diamonds). (C) Empty vector (vector) cells, 7A 1-2 cells, 7A 1-4 cells, 7B 1-3 cells, and 7B 2-2 cells were plated and treated with romidepsin (10 ng/ml), KD-5170 (10 µM), largazole (250 nM), NCH-51 (25 µM), or OKI-005 (500 nM) for 24 h after which protein was extracted and subjected to PAGE. Proteins were transferred to nitrocellulose membranes and blots were incubated with antibodies to pan acetylated histone H3 (AcH3), total histone H3 (total H3), p21 or actin. Results from one of three independent experiments is shown. (D) Empty vector (vector) cells, 7A 1-2 cells, 7A 1-4 cells, 7B 1-3 cells, and 7B 2-2 cells were plated and treated with romidepsin (10 ng/ml), KD-5170 (10 µM), largazole (250 nM), NCH-51 (25 µM), or OKI-005 (500 nM) for 24 h after which cell cycle analysis was performed. The percentage of cells in each phase was determined using ModFit LT v 5.0. The graph was generated from data compiled from 3 independent experiments. Error bars indicate standard deviation.

We next treated HEK-293 cells transfected with empty vector (Vector) or one of the METTL7A (7A 1-2) and METTL7B (7B 2-2) clones with romidepsin, KD-5170, largazole, NCH-51, or OKI-005 for 24 h and examined changes in histone acetylation and p21 expression. In agreement with cytotoxicity assays, cells expressing METTL7B were not resistant to the effects of romidepsin and displayed increased levels of acetylated histone H3 and p21, while cells expressing METTL7A were resistant to the effects of all the thiol-based HDACis (Fig. 5C).

Cell cycle analysis was also performed on the transfected cells after treatment with romidepsin, KD-5170, largazole, NCH-51, or OKI-005 for 24 h. As was observed with MCF-7 cells, all thiol-based HDACs induced cell cycle changes in empty-vector transfected cells, causing a decrease in the number of cells in the G_1_/G_0_ and S phases and an increase in the number of cells in the G_2_/M phase (Supplemental Fig. S4). In contrast, no significant cell cycle changes were observed in cells transfected to express METTL7A. In the METTL7B clones, while no significant cell cycle changes were observed with a majority of the thiol-based HDACs, we did note significant cell cycle perturbations in cells treated with romidepsin and KD-5170 (Fig. 5D).

### Evidence of methylated romidepsin in culture media

If METTL7A can methylate romidepsin, we rationalized that we should observe methylated romidepsin in the culture medium of MCF-7 DpVp300 cells incubated with the drug. Using liquid-chromatography coupled to tandem-mass spectrometry (LC-MS/MS) we observed high levels of dimethylated romidepsin in the media of MCF-7 DpVp300 cells (Fig. 6A), whereas much lower levels were observed in the media of MCF-7 cells and the two METTL7A knockout clones, supporting the hypothesis that METTL7A can methylate and inactivate romidepsin.

**Figure 6.**
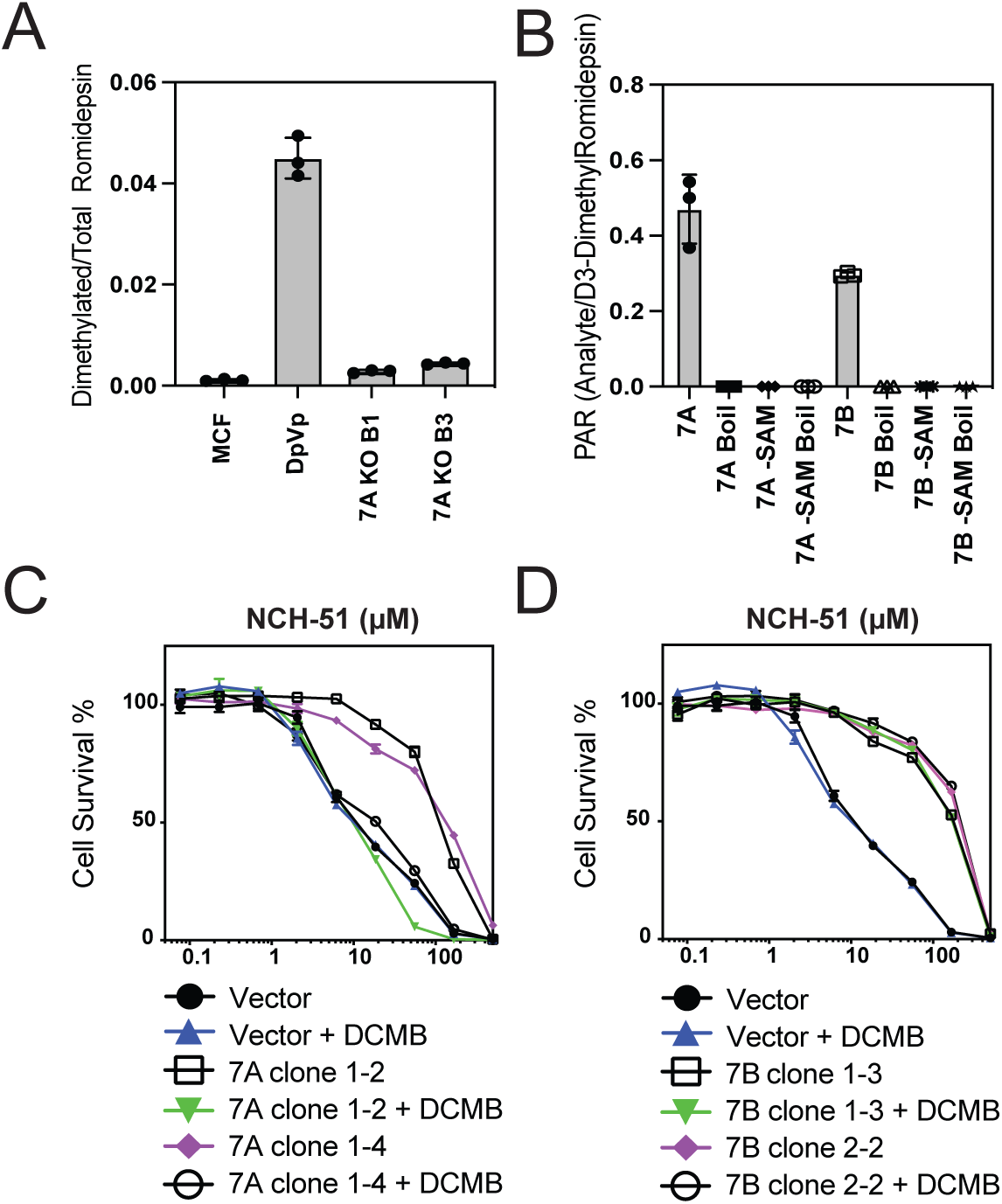
METTL7A methylation of romidepsin *in vitro*. (A) Measurement of dimethylated romidepsin by LC-MS/MS in cell culture media following 24 h of incubation with MCF-7, MCF-7 DpVp300, 7A KO B1 and 7A KO B3 cells. (B) Measurement of methylated romidepsin with METTL7A and METTL7B in a cell-free system. Methylation reactions were carried out with recombinant enzyme and the amount of methylated product was measured by LC-MS/MS. Omission of the methyl donor SAM, and boiling the enzyme were used as controls. (C) Three-day cytotoxicity assays were performed with NCH-51 on empty vector (vector) cells (black circles), empty vector cells with DCMB (blue triangles) 7A 1-2 cells (open black squares) 7A 1-2 cells with DCMB (inverted green triangles) 7A 1-4 cells (purple diamonds), and 7A 1-2 cells with DCMB (open black circles). (D) Three-day cytotoxicity assays were performed with NCH-51 on empty vector (vector) cells (black circles), empty vector cells with DCMB (blue triangles) 7B 1-3 cells (open black squares) 7B 1-3 cells with DCMB (inverted green triangles) 7B 2-2 cells (purple diamonds), and 7B 2-2 cells with DCMB (open black circles).

The ability of METTLA and METTL7B to methylate romidepsin was also examined in a cell-free system. Recombinant METTL7A and METTL7B were incubated with reduced romidepsin for 2 hours and mass spectrometry performed to quantify any monomethylate reduced romidepsin. Both METTL7A and METTL7B were found to methylate reduced romidepsin (Fig. 6B). In contrast, no methylated product, or trace amounts was noted when the reactions were performed in the absence of the methyl donor S-adenosyl-L-methionine (SAM), or when the enzyme was pre-treated with boiling. Together, these results confirm the methyltransferase activity of these enzymes using romidepsin as a substrate.

The phenylethanolamine-N-methyltransferase inhibitor 2,3-Dichloro-α-methylbenzylamine hydrochloride (DCMB) was recently reported to selectively inhibit the methyltransferase activity of METTL7A but not METTL7B [30]. We therefore conducted cytotoxicity assays in cells over expressing METTL7A or METTL7B combining DCMB and NCH-51, since NCH-51 was found to be a substrate of both METTL7A and METTL7B. Using a non-toxic concentration of DCMB we found that DCMB completely reversed resistance to NCH-51 mediated by METTL7A but not METTL7B. Inhibitors of other methyltransferases could also be potential inhibitors of METTL7A and METTL7B and may reverse resistance to thiol-based HDACis.

### Tissue expression of METTL7A and METTL7B

While our work with resistant cells implicates METTL7A in resistance to HDACis, its physiological role in development and homeostasis remains to be determined. To understand the localization of METTL7A and METTL7B within tissues, we performed immunohistochemistry on normal tissue microarrays. Both METTL7A and METTL7B were found to be highly expressed in hepatocytes (Fig 7A and 7B, respectively) and distinct segments of the nephron (Fig. 7C and 7D), while METTL7B was found in the intestinal enterocytes (Fig. 7E and 7F) and METTL7A was found in breast epithelium (Fig. 7G and 7H). METLL7A is also highly expressed in glial cells, certain nephron segments, and Leydig cells of the testes, while METTL7B is highly expressed in enterocytes, neurons, and certain nephron segments. Modest expression of both METTL7A and METTL7B is also found in cardiomyocytes, gastric mucosa, and subcutaneous adipocytes (Sup. Table 5). The localization pattern of METTL7A and METTL7B is consistent with a role in detoxification, which might have treatment implications for drugs in this class.

**Figure 7:**
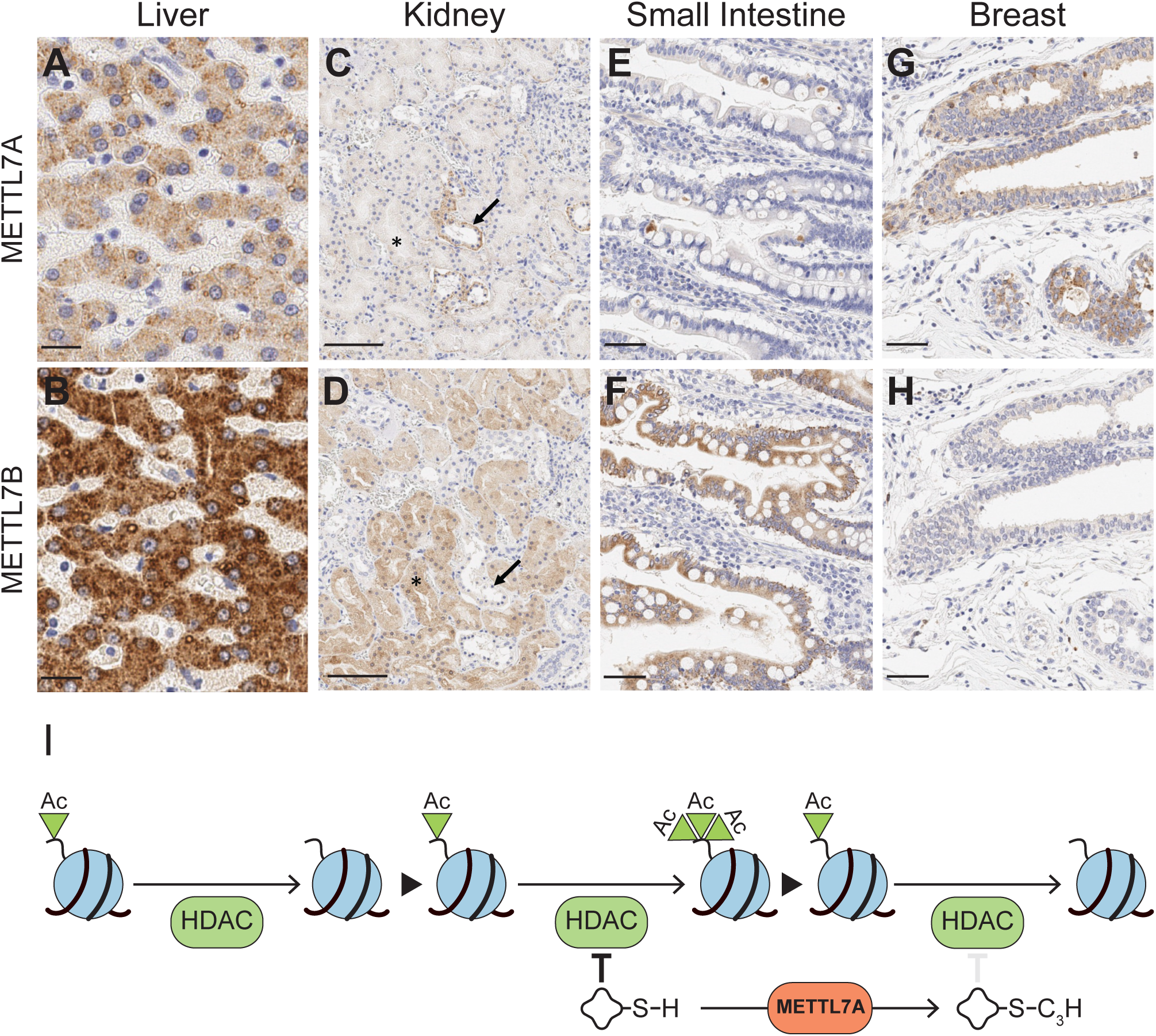
Normal Tissue Expression and proposed model. Normal tissue expression of METTL7A and METTL7B. A strong cytoplasmic pattern is observed in hepatocytes (A and B), distinct regions of the nephron (C and D), small intestinal enterocytes (E and F), and breast epithelium (G and H). Arrows and asterisks indicate distinct METTL7A and METTL7B expression patterns in renal tubules on serial sections. Scale bars represent 20 µm (A and B), 100 µm (C and D), and 50 µm (E, F, G, and H). (I) Proposed model of drug resistance through activity of METTL7A.

## DISCUSSION

Precise control of gene expression through acetylation and deacetylation of lysine residues within the tails of histones is important for timely expression of genes, and this epigenetic regulation is frequently dysregulated in cancers (reviewed in [31]). HDACis are a novel class of chemotherapeutic drugs that modulate the acetylation levels of histones by coordinating with Zn^2+^ in the catalytic pocket of HDACs, thus inhibiting enzyme activity. Reduced HDAC activity results in hyperacetylation of substrates and induces downstream cell cycle arrest or apoptosis. Vorinostat and romidepsin were the first FDA-approved HDACis for the treatment of T-cell lymphomas [5–7, 32]. Despite dramatic responses in some patients, romidepsin is not always effective and resistance can develop after responses. Furthermore, the activity of HDACis in solid tumors has been limited, suggesting intrinsic mechanisms of resistance. While elevated levels of the efflux pump P-gp are associated with *in vitro* resistance to romidepsin, this mechanism does not translate to the clinic [17]. Thus, we set out to identify mechanisms of resistance to HDACis. Here, we identify overexpression of the putative methyltransferases METTL7A and METTL7B as a mechanism of drug resistance to thiol-based HDACis (Fig. 7I).

In this study, we identified overexpression of METTL7A as a mechanism of romidepsin resistance in two different cell models. In both models, we uncovered a mechanism of resistance that is independent of P-gp function and acts upstream of the deacetylation process. Expression of the methyltransferase METTL7A is necessary for resistance, and expression of METTL7A in naïve cells can drive resistance to thiol-containing HDACis. METTL7A-dependent resistance to romidepsin correlated with the presence of methylated drug in cell media, and the ability of METTL7A to drive resistance to thiol-containing HDACis can be blocked by the methyltransferase inhibitor DCMB. Finally, METTL7A can methylate romidesin *in vitro*. Our data supports the model whereby exposure of cells to romidepsin selects for upregulation of the methyltransferase METTL7A. Once expressed, METTL7A methylates the zinc-coordinating thiol to prevents the drug from inhibiting HDAC, rendering the cells resistant to this class of therapeutic agents. We have noted that the cells selected for high levels of resistance to romidepsin are also resistant, albeit to a lesser extent, to other HDACis that do not contain thiol groups. This suggests that one or more factors in the resistant cells promote more general resistance to HDACis. Whether these are mediated by some of the other molecular changes we have observed in these cells remains to be determined. Several mechanisms might be involved in METTL7A upregulation. As previously described [33], we observe extensive editing of the 3′-UTR of METTL7A in both of our resistance models (Supplemental Fig. S2D), suggesting that post-transcriptional regulation may play a role in the up-regulation of METTL7A. We also observe changes in chromatin accessibility at the METTL7A locus that support up-regulation of METTL7A at the transcriptional level. The upregulation in METTL7A transcription in the DpVp cells could be driven by changes in transcriptional networks, changes in DNA methylation at CpG sites in the METTL7A gene body, or both. In thyroid cancer, expression of METTL7A has been shown to be regulated by methylation of a CpG island at exon 2 of *METTL7A* [34]. Since HDACis have been shown to affect DNA methylation [35–37], one hypothesis is that exposure to HDACis results in a reduction of DNA methylation at the METTL7A locus that allows for the expression of the METTL7A mRNA.

Few studies have linked expression of METTL7A or METTL7B to drug resistance. Jun and colleagues identified METTL7A as a mechanism of resistance in methotrexate resistant choriocarcinoma cells [38]. However, when we performed cytotoxicity assays with methotrexate on MCF-7 and DpVp300 cells, we did not find increased resistance in the DpVp300 line. Additionally, we note that methotrexate does not contain thiol groups. Thus, other potential factors besides METTL7A may be involved in methotrexate resistance. A role for METTL7B in resistance to tyrosine kinase inhibitors has been suggested by Song and colleagues who noted increased expression of METTL7B in lung adenocarcinoma cell lines that were resistant to the tyrosine kinase inhibitors (TKIs) gefitinib or osimertinib [39]. In their report, METTL7B was not found to directly interact with the TKIs, rather METTL7B overexpression was associated with increased antioxidant capacity and reactive oxygen species (ROS) scavenging [39].

While two previous studies suggested METTL7A methylates RNA [40] and DNA [41], our studies, in line with the work of Totah and colleagues on both METTL7A [30] and the close paralog METTL7B [25], suggest that METTL7A directly methylates thiol groups on small molecules in the cell.

While we observed that both METTL7A and METTL7B were able to methylate romidepsin in a cell-free system, experiments with our cell line models suggests that the two paralogs have different roles in drug resistance, with METTL7A being the relevant driver of resistance to romidepsin. Several reasons can explain this discrepancy and warrant further investigation. Differences in reaction site preferences, post-translational modifications, binding partners, or sub-cellular localization between the two enzymes could results in differences in activity toward romidepsin.

While our work reveals a role for METTL7A and METTL7B in drug resistance, the function of these methyltransferases during development and a physiological role for them remains to be elucidated. One of the murine homologs of METTL7A, Mettl7a1, is transiently and significantly upregulated during reprogramming of mouse embryonic fibroblasts (MEFs) to induce endoderm progenitors [42]. Addition of Mettl7a1 to the reprogramming cocktail enhanced conversion of MEFs into induced endoderm progenitors by promoting reprogramming efficiency. METTL7A also plays a role in osteogenic differentiation by mediating the effects of glucose on cell survival and osteogenic induction [41]. Lastly, a role in hydrogen sulfide homeostasis has also been proposed for both METTL7A and METTL7B [25, 30]. Both paralogs are expressed in the stomach, kidney, and liver which may suggest a detoxifying role for these proteins under normal physiological conditions.

The data presented here show that METTL7A has the ability to inactivate thiol-containing HDAC inhibitors before they reach their targets. It is possible that this mechanism of resistance, despite the potential to reduce effective drug concentrations in cancer cells, reduces the systemic effects of the drugs given to patients. The key challenge in oncopharmacology is that we do not always know if a drug reached its target. How much the presence of METTL7A is responsible for the inability to translate the unique activity seen in T-cell lymphomas to solid tumors remains to be investigated ([43–46]). We can further hypothesize a role in the emergence of resistance in T-cell lymphomas, and this remains to be investigated as well. Together, these findings are a reminder of just how much remains to be learned and solved regarding how cancers become resistant to treatment.

## ACKNOWLEDGEMENTS

We thank members of the Batista, Gottesman and Arda Labs for discussion and critical reading of the manuscript. The authors thank the Center for Cancer Research (CCR) Genomics Core in Bethesda, Maryland, as well as the CCR Sequencing Facility in Frederick, Maryland, for help with high-throughput sequencing. This work utilized the computational resources of the NIH HPC Biowulf cluster (http://hpc.nih.gov). We appreciate the editorial assistance of George Leiman. The views expressed in this article are those of authors and may not reflect the official policy or position of the National Institute of Health, Department of the Army, Department of Defense. Mention of trade names, commercial products, or organizations imply endorsement by the U.S. Government. Supported by the Intramural Research Program at the National Cancer Institute (NCI) of the National Institutes of Health. A.K.T. is supported by Department of Defense – BCRP Level 2 award (1W81XWH-21-1-0053). C.M.F. is partially supported by a Postdoctoral Fellowship from the American Cancer Society (PF-19-157-01-RMC). Authors’ Discosures: Anthony D. Piscopio is CEO and president of OnKure, Inc

## HIGH-THROUGHPUT SEQUENCING DATA AND CODE AVAILABILITY

RNA-seq and ATAC-seq data were deposited in the Gene Expression Omnibus database GSE217130. Code and scripts used to generate data and plots are available at https://github.com/BatistaLab/METTL7A

## MATERIALS AND METHODS

### Chemicals

Methotrexate, 5-fluoruracil, teniposide, apicidin, trichostatin a, and topotecan were obtained from Sigma Aldrich (St. Louis, MO). Romidepsin was obtained from Selleck Chem (Houston, TX). Belinostat, mocetinostat, and panobinostat were purchased from ChemieTek (Indiannapolis, IN). Dacinostat, resminostat, pracinostat, chidamide, vorinostat, entinostat, NCH-51, and KD 5170 were obtained from Cayman Chemical (Ann Arbor, MI). Rocilinostat was purchased from ApexBio Technology (Houston, TX). PCI-34051 was from Tocris (Minneapolis, MN). Largazole and OKI-005 were synthesized in-house by OnKure.

### Cells

MCF-7 breast cancer cells were selected in increasing concentrations of romidepsin to generate the MCF-7 DpVp300 cell line that is maintained in 300 ng/ml romidepsin with 5 µg/ml verapamil. The HUT78 cell line was purchased from ATCC (Manassas, VA) and the romidepsin-resistant HUT78 DpVp50 and DpP75 cell lines have been previously described [18]. All cells were cultured in RPMI-1640 supplemented with 10% FCS and Pen/Strep. HEK293 cells were also from ATCC and were maintained in Eagle’s minimum essential medium with 10% FCS and Pen/Strep. HEK293 cells were transfected with empty pcDNA3.1 vector, or vector containing full-length *METTL7A* or *METTL7B* with a C-terminus FLAG tag (all vectors from Genscript, Piscataway, NJ) using Lipofectamine 2000 (Thermo Fisher Scientific, Waltham, MA) according to the manufacturer’s instructions. Clones were selected and were maintained in 1 mg/mL G418. CRISPR-mediated knockout of METTL7A from MCF-7 DpVp300 cells was achieved by co-transfecting knockout and homology directed repair vectors for METTL7A (obtained from Santa Cruz Biotechnology, Dallas, TX) using Lipofectamine 2000 and subsequent selection with puromycin (3 µg/ml). Clones were then isolated and characterized for loss of METTL7A protein via immunoblot.

### Cytotoxicity assays

Cells were plated at a density of 5000 cells/well in opaque, white 96-well plates and allowed to attach overnight. The next day drugs were added, and plates were incubated for 3 days after which CellTiterGlo (Promega, Madison, WI) was added according to the manufacturer’s instructions. Plates were read on a Tecan Infinite M200 Pro (Tecan Systems, Inc., San José, CA).

### Immunoblot analysis

Whole cell lysates were obtained by harvesting cells in RIPA buffer and sonicating samples in an inverted cup horn sonicator for 3 bursts of 30 sec, after which the samples were centrifuged at 10000 RPM for 10 min and soluble proteins were removed. When examining histone acetylation, RIPA buffer additionally contained 500 nM trichostatin A to inhibit histone deacetylases after lysing cells. Protein lysates (approx. 15-20 µg) were separated by PAGE and transferred to nitrocellulose membranes that were subsequently probed with antibodies against total histone H3 (cat# 05-499, Millipore-Sigma, Burlington, MA), pan-Acetyl histone H3 (cat# 06-599, Millipore-Sigma), METTL7A, METTL7B, beta-actin, (generated in mouse cat# 3700, or rabbit cat# 4970, Cell Signaling Technology, Danvers, MA) and p21 (cat# 2946, Cell Signaling). Anti-FLAG antibody (cat# F1804-200UG) was from also from Millipore-Sigma.

### Flow cytometry

To determine surface expression of P-glycoprotein, trypsinized cells were incubated with phycoerythrin-labeled UIC2 antibody or phycoerythrin-labeled isotype control IgG2a kappa (both from ThermoFisher, Grand Island, NY) for 20 min at room temperature in 2% bovine serum albumin/PBS. Cell fluorescence was measured with a FACSCanto flow cytometer (BD Biosciences, San José, CA) and data analysis was performed using FloJo v10.4.2 (FlowJo LLC, Ashland OR). At least 10,000 events were collected per sample. For cell cycle analysis, cells were plated and allowed to attach overnight before being treated with HDACIs for 24 h. Cells were then harvested by trypsinization, centrifuged, and the supernatant removed. Cell pellets were resuspended in equal volumes of staining solution (0.05 mg/ml propidium iodide and 0.1% Triton X) and RNAse A solution (200 U/mL RNAse A in deionized water). Cells were stained for 20 min and were subsequently read on a FACSCanto Flow Cytometer with at least 10,000 events collected per sample. The percentage of cells in each phase of the cell cycle was determined by Modfit LT for Mac version 5.0 (Verity Software House, Topsham, ME). Statistically significant changes in the percent of cells in each phase of the cell cycle was determined by a one-way ANOVA test with a correction for multiple comparisons.

### RNA sequencing analysis

RNA was extracted from the MCF-7, MCF-7 DpVp300, HuT78, HuT78 DpVp50 and HuT78 DpP50 cell lines using the RNeasy kit from Qiagen (Germantown, MD). RNA sample integrity and quantity for three biological replicates were assessed by the Center for Cancer Research Genomics Core using RNA ScreenTape Analysis (Agilent). All RNA samples had RNA Integrity Numbers of (> 8.0). RNA libraries were generated using a TruSeq Stranded mRNA library prep kit (Illumnia) and paired-end RNA sequencing (2 x 75) was performed by the CCR Sequencing Facility and Genomics Technology Core on a NextSeq 500 Sequencing System. Read quality analysis was performed using FASTQC. Reads of the samples were trimmed for adapters and low-quality bases using Cutadapt (version 2.3) before alignment with the hg38 reference genome using Spliced Transcripts Alignment to a Reference (version 2.7.0f). The number of reads mapping to each transcript was determined with HTseq-count (version 0.9.1). Differential expression was determined using DESeq2 (version 1.26.0) within the R Studio (version 3.6.3).

### Omni-ATAC-seq assay

#### Library Generation

Approximately 20,000 cells per sample were used for omni ATAC-seq assay following the protocol described in Corces et al. [47] with minor modifications. For each cell line, two biological replicates were included for control and resistant lines, respectively. To isolate the nuclei, cells were resuspended in cold ATAC-Resuspension Buffer (RSB) containing 0.1% NP40, 0.1% Tween-20, and 0.01% Digitonin, and incubated on ice for 3 minutes. The lysis was washed out by addition of 1 mL of RSB containing 0.1% Tween-20 and mixed by inverting tubes. The nuclei were then pelleted at 500RCF for 10 minutes at 4C. For transposition, each nuclei pellet was resuspended in 50 µL of transposition mix (25 µl 2x TD buffer, 2.5 µl transposase (100 nM final), 16.5 µl PBS, 0.5 µl 1% digitonin, 0.5 µl 10% Tween-20, 5 µl H_2_O) and incubated at 37C for 30 minutes in a thermomixer with 1000 rpm shaking. After transposition, reactions were stopped by adding EDTA to a final concentration of 40 mM. Transposed DNA fragments were purified using Zymo DNA Clean and Concentrator-5 Kit (cat# D4014) and PCR amplified for 5 cycles. Each PCR reaction contained 2.5 µl of 25 µM i5 primer, 2.5 µl of 25 µM i7 primer, 25 µl 2x NEBNext high-fidelity 2x master mix (NEB # M0541), and 20 µl transposed/cleaned-up DNA sample. PCR conditions were as follows: 72°C for 5 min, 98°C for 30 sec, followed by 5 cycles of [98°C for 10 sec, 63°C for 30 sec, 72°C for 1 min] and then kept at 4°C. PCR reactions were purified using the Zymo kit. The purified ATAC libraries were quantified on an Aligent TapeStation using D5000 high sensitivity tapes and mixed at an equal molar ratio. The library pool then was size-selected for 100-1000 bp on a 1% agarose gel. The final library pool was sequenced on Illumina NextSeq2000 for a 100 bp pair-ended run to obtain 100 million reads per sample.

#### ATAC-seq Analysis

FASTQ files were processed using the PEPATAC (v 2.0.0) pipeline [48] against the hg38 build of the human genome. Briefly, reads were trimmed of adaptors using Skewer and aligned to hg38 using Bowtie2. Duplicates were removed using Samblaster. Peaks were called using MACS2. Differently accessible regions were determined using DESeq2 (version 1.26.0) within R Studio (version 3.6.3). *De novo* and known motif enrichment analysis was performed with HOMER [49]. Changes in TF occupancy between the two cell states was performed with the BaGFoot algorithm [50].

### Chromatin extraction and MNase+ExoIII-seq assay

Cells were harvested at 80% confluency using a cell scraper, centrifuged at 125g for 5 min and washed once with cold phosphate-buffered saline. Cells were then resuspended in cold RSB buffer (10 mM Tris-HCl, pH 8.0, 10 mM NaCl, 3 mM MgCl_2_) containing 0.5% Nonidet P-40, 1 mM PMSF and put on ice. The cell suspensions were homogenized by 20 strokes of pestle B in a Dounce homogenizer over 30 min on ice. Release nuclei were centrifuged in Sorvall X Pro centrifuge at 150g, 4°C for 5 min, nuclei pellet was resuspended in cold RSB buffer supplemented with 1 mM PMSF.

Before digestion, 1×10 ^6^ nuclei in RSB buffer with 1 mM PMSF were supplemented with 2 mM CaCl_2_ and kept at 37°C for 5 min. 5 units of Micrococcal nuclease (Nuclease S7 Micrococcal Nuclease, cat# 10107921001; Roche Applied Science) with 200 units ExoIII (cat# M0206S; New England Biolabs) were added to each nuclei aliquot and incubated for 10 min at 37°C. The reaction was stopped by adding EDTA to a final concentration of 5 mM. Digested nuclei were centrifuged at 4,600g in a Sorvall Legend Micro tabletop centrifuge, supernatant S1 was removed and discarded. Pellet was resuspended in cold TE buffer, (10 mM Tris-HCl pH 8.0, 1 mM EDTA), gently pipetted and incubated on ice for 10 min. After centrifugation supernatant S2 was collected, pelleted material was resuspended in TE buffer, and after third centrifugation supernatant S3 was collected. Supernatants 2 and 3 were analyzed on 1% agarose gel in TAE buffer to confirm the release of DNA fragments of various length, then combined and treated with 1% SDS and 20 µg of proteinase K (New England Biolabs, #P8107S). The reaction was incubated for 1 h at 55°C. DNA was extracted with phenol-chloroform-isoamyl alcohol, precipitated in 2.5 volumes of 100% alcohol supplemented with 50 mM sodium-acetate and left at -80°C for overnight. Samples were centrifuged at 17,000g at a tabletop centrifuge, pellet was rinsed in 70% alcohol, air dried and dissolved in TE buffer (10 mM Tris-HCl, pH 8.0, 0.2 mM EDTA). Total DNA was sequenced on Illumina NextSeq2000 for a 100 bp pair-ended run to obtain 300 million reads per sample.

### LC-MS to detect methylated romidepsin in culture medium

Media was centrifuged 5000 × g 15 minutes to pellet any insoluble material and 50 µL of supernatant removed for mass spectrometry analysis. For each, 10 10 µL was injected on an Exion liquid chromatography system (SCIEX, Framingham, MA) coupled inline to an X500B QTOF mass spectrometer (SCIEX, Framingham, MA) in-line with an Exion liquid chromatography system (SCIEX, Framingham, MA). Each sample was separated on a 2.1 x 50 mm Aeris 3.6 µm WIDEPORE XB-C8 200 Å column (Phenomenex, Torrance, CA) at 0.3 mL/min using a 4 min linear gradient from 98% A (0.1% formic acid) to 40% B (acetonitrile with 0.1% formic acid). Data was acquired in positive mode over a mass range of 100-800 m/z with data dependent acquisition MS/MS performed over the mass range of 50-750 m/z. Relative quantitation was performed from extracted ion chromatogram peak areas using SCIEX OS software (SCIEX, Framingham, MA).

### Reduction of romidepsin

Romidepsin was added to 1 mL of deoxygenated 50 mM Tris-HCl buffer pH 7.5 with 10 mM TCEP. The final romidepsin concentration was 1 mM and was incubated at 37 °C for 1.5 h in the TCEP-containing, buffered solution. At 1.5 h the romidepsin, now almost completely reduced, was extracted with 1X ethyl acetate. Briefly, 1mL of ethyl acetate was added to the 1 mL solution of, now reduced, romidepsin. The entire solution was vortexed, centrifuged briefly at 5000 x g, and then incubated at -80 °C to freeze the aqueous components. The top layer of ethyl acetate does not freeze and was transferred to a new vial and dried with a slow stream of nitrogen gas. The dried reduced romidepsin was then reconstituted with acetonitrile.

### Synthesis of (D3)2-diemthyl-romidepsin

We determined that, when reduced with TCEP as previously described, romidepsin was almost completely converted to the reduced form. We therefore assumed that when reconstituted with acetonitrile, the solution contains approx. 1 µmol of reduced romidepsin. All reagents are added based on this estimated yield of reduced romidepsin. To a solution of acetonitrile containing 1 µmol of reduced romidepsin dissolved in acetonitrile, 10 µmol of N,N-Diisopropylethylamine (DIPEA) and 5 umol of D3-iodomethane were added. The solution was incubated at room temperature for 2 h before adding a small volume of water to quench the remaining iodomethane. The solution was then dried down with a slow stream of nitrogen gas. The dried D3-dimethyl-romidepsin was then reconstituted with acetonitrile and stored at -80 °C.

### Romidepsin methylation activity with purified enzyme

Purified N-GST-METTL7A and -METTL7B were individually diluted to 0.2 mg/mL in KPi reaction buffer (50 mM KPi pH 7.0, 20% glycerol, 150 mM NaCl, 10 mM CHAPS). The diluted purified thiol methyltransferases were then added 1:2 to a solution of 9 mg/mL 1,2-Dimyristoyl-sn-glycero-3-phospho-rac-(1-glycerol) sodium salt (DMPG) dissolved in KPi reaction buffer. The final protein concentration was 0.07 mg/mL for each protein.

The solutions of diluted protein were incubated on ice for 30 mins. Small volumes of diluted METTL7A and METTL7B were then aliquoted into separate vials and boiled for 5 min to inactivate the enzymes. A small volume of reduced romidepsin was deposited into relevant wells of a 96-well reaction plate. The acetonitrile solvent was removed by placing the plate in a vacuum chamber and pulling a vacuum for several minutes. The diluted proteins were added to the plate wells containing dried reduced romidepsin.

Vehicle control was then added to relevant wells containing the diluted thiol methyltransferases. The well plate was allowed to shake at 625 rpm on a plate shaker for 5 min. Finally, SAM or a vehicle control of liquid chromatography-grade water was added to the appropriate wells to initiate the reaction. The final concentration of protein was 0.06 mg/mL and the final concentration of SAM was 500 µM. The estimated final concentration of reduced romidepsin was approx. 500 µM. After SAM was added to initiate the reaction, the samples were incubated at 37 °C for 2 h. After 2 h, ice-cold acetonitrile containing the (D3)2-dimethyl-romidepsin internal standard was added to precipitate salts and proteins at a ratio of 3:7 (reaction volume:MeCN). The plate was centrifuged at 5000 x g for 20 min and then the supernatant from each well was transferred to a new well in a 96-well plate for LC-MS analysis.

Samples were analyzed with a Waters Xevo TQS mass spectrometer paired with a Waters Acquity LC using a 2.1 × 50 mm Waters 1.7 µm C18 column and 0.2% acetic acid in water (Buffer A) and 0.2% acetic acid in MeCN (Buffer B). Consistent elution of romidepsin analytes was achieved with the following gradient: solvent B held at 5% from 0 to 0.15 min, then increased to 95% from 0.15 to 5 min, followed by re-equilibration to the starting conditions for another 2 min for a total run time of 7 min. Flow rate was held constant at 0.35 mL/min. The transition used to monitor for the early monomethyl metabolites was 557.74> 255.

### Immunohistochemistry

Serial sections from normal tissue microarrays (BCN921 and BN1021) were obtained from tissuearray.com (Derwood, MD). Slides were immunolabeled for METTLA and METTLB according to the conditions in Supplemental Table S6. Serial sections were also stained with a negative control protocol in which primary antibody was replaced by nonclonal antibody from the same species and of the same isotypes (isotype controls). Positive controls included cell pellet slides with known expression of METTL7A (7A 1-2, 7A 1-4) and METTL7B (7B 1-3, 7B 2-2). Slides were analyzed and each tissue was manually evaluated for positive staining along with negative controls; positive cell types were listed and assigned a semi-quantitative grade for positivity (minimal, mild, moderate, or marked; 1+, 2+, 3+, 4+). Slides were also digitalized and analyzed using QuPath (v 0.3.2) [51]. Digital image analysis included identifying stain vectors, de-arraying tissue microarrays, defining regions of interest, cell detection, and quantifying chromogenic immunohistochemistry staining. Following automated TMA grid detection, manual pathologist review was performed to exclude spots or regions of spots that were non-assessable, non-tumor containing, or obscured by artifact.

## Figure Legends

**Supplemental Fig. S1.**
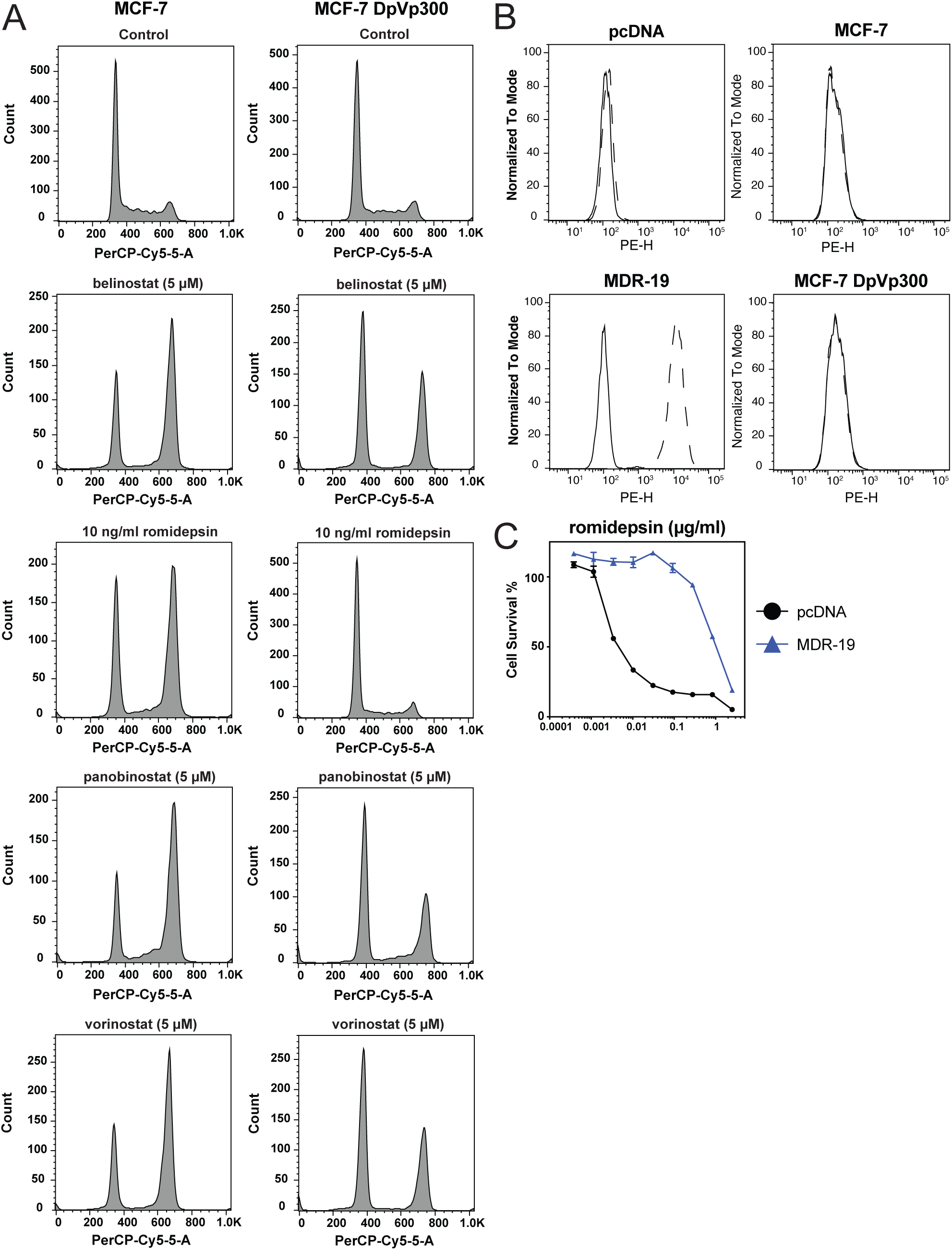
MCF-7 DpVp300 are selectively resistant to romidepsin and do not overexpress P-gp. (A) MCF-7 and MCF-7 DpVp300 cells were plated and treated with the noted concentrations of HDACI for 24 h, after which cells were harvested and cell cycle analysis was performed as detailed in the materials and methods. (B) MCF-7, MCF-7 DpVp300, pcDNA, and MDR-19 cells were trypsinized and incubated with isotype control antibody (solid line) or UIC-2 antibody (dashed line) and fluorescence was measured on a FACSCanto flow cytometer. (C) Three-day cytotoxicity assays were performed with romidepsin on pcDNA cells (black circles) and MDR-19 (blue triangles). Results from one of three experiments is shown.

**Supplemental Fig. S2.**
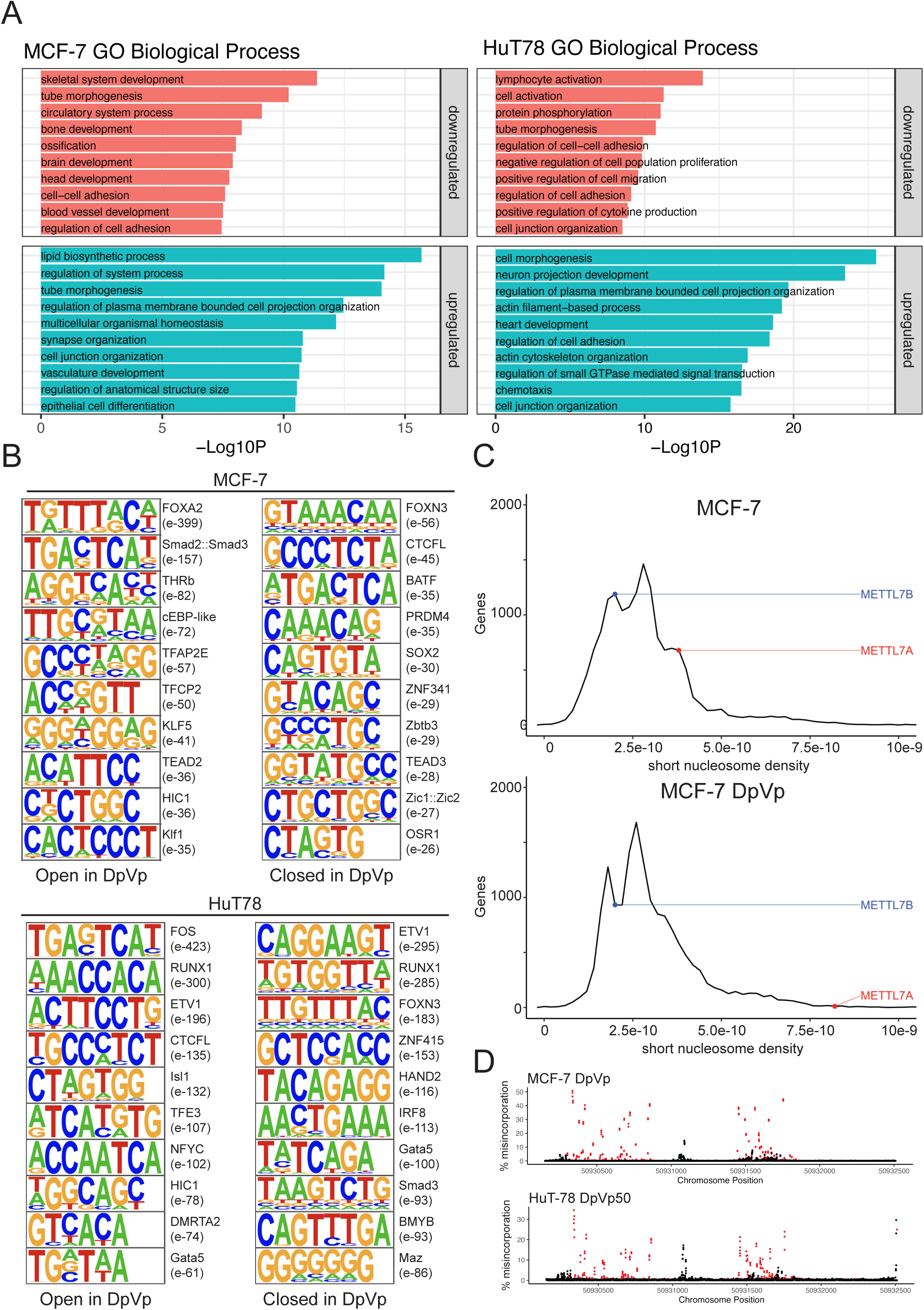
RNA Seq and ATAC Seq analysis of MCF-7, MCF-7 DpVp300, HuT78 and HuT78 DpVp50 cells. (A) Gene ontology (GO)-Term analysis of up- and down-regulated genes from RNA-seq analysis in the MCF-7 and HuT-78 romidpesin resistance models. (B) *De novo* motif analysis of open DORs or closed DORs in the MCF-7 and Hut-78 romidepsin resistance models. (C) Histogram representation of short nucleosome (80-140bp) density in MCF-7 and MCF-7 DpVp300 cells. METTL7B is shown as example of a gene that does not change. (D) Histogram representing misincorporation frequency in RNA sequencing experiments in MCF-7 DpVp300 and HuT78 DpVp50 cells. Red represents A to G changes, black all other changes.

**Supplemental Fig. S3.**
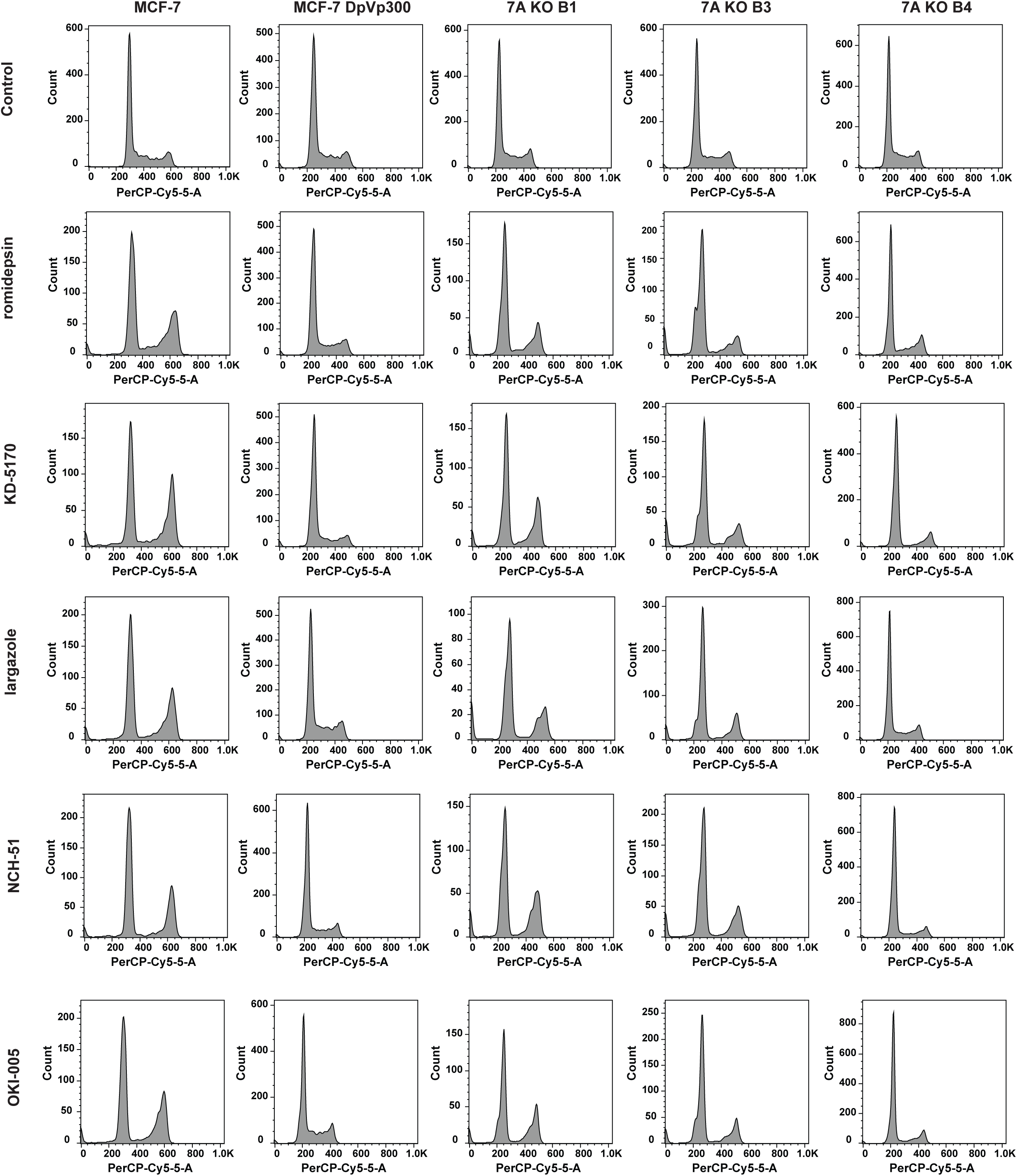
Thiol-based HDACIs induce cell cycle changes after deletion of METTL7A in MCF-7 DpVp300 cells. MCF-7, MCF-7 DpVp300, 7A KO B1, 7A KO B3 and 7A KO B4 cells were plated and treated with the noted concentrations of HDACi for 24 h, after which cells were harvested and cell cycle analysis was performed as detailed in Materials and Methods.

**Supplemental Fig. S4.**
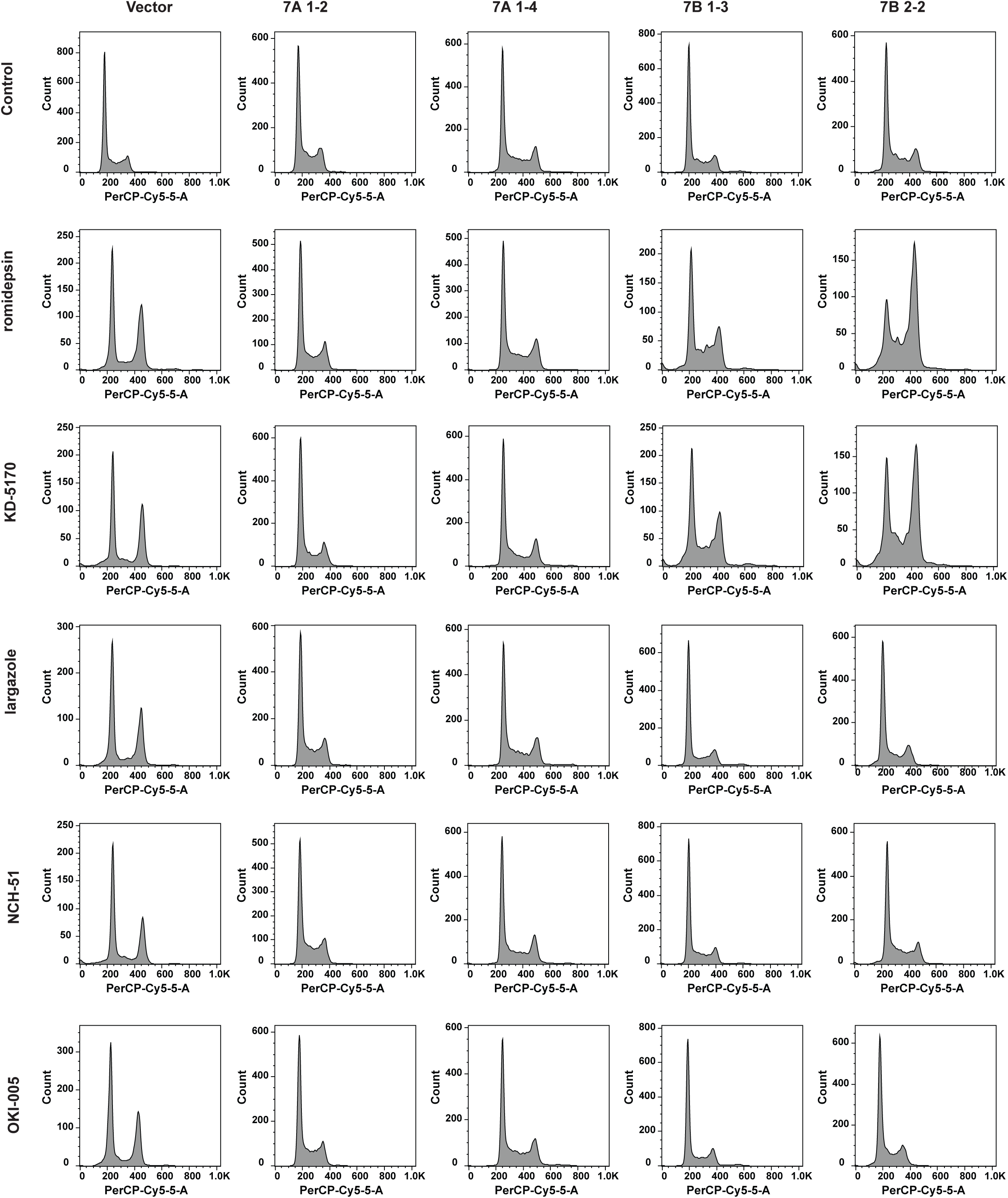
METTL7A overexpression prevents cell cycle changes mediated by thiol-based HDACis Vector, 7A 1-2, 7A 1-4, 7B 1-3 and 7B 2-2 cells were plated and treated with the noted concentrations of HDACI for 24 h, after which cells were harvested and cell cycle analysis was performed as detailed in Materials and Methods.

**Supplemental Table S1.**
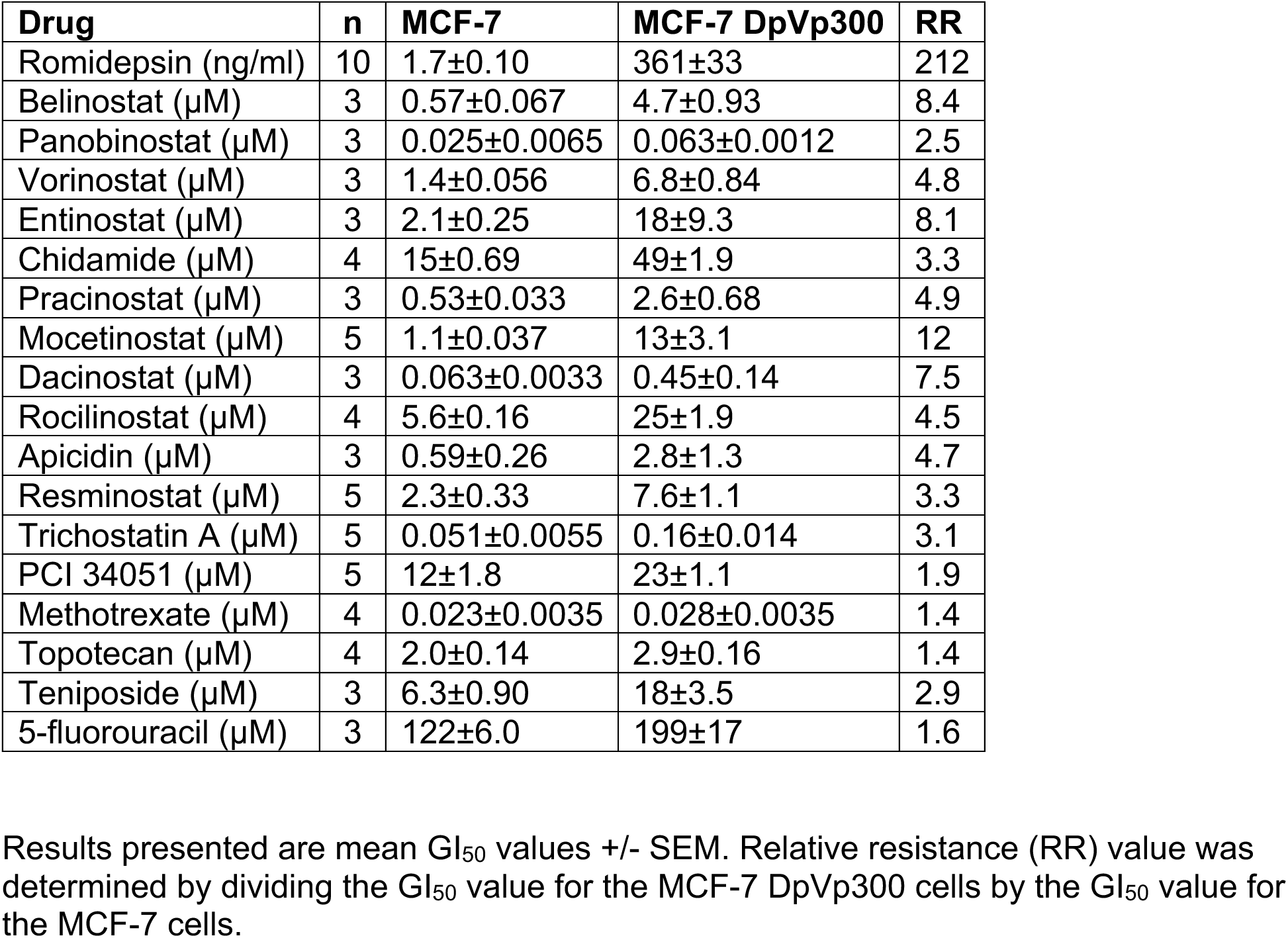
Cross resistance profile of MCF-7 DpVp300 cells

**Supplemental Table S2.** Cross resistance profile of MCF-7 DpVp300 cells with thiol-based HDACIs

| Drug | n | MCF-7 | DpVp300 | RR |
| --- | --- | --- | --- | --- |
| KD5170 | 3 | 4.9±1.3 | 173±11 | 35 |
| Largazole | 3 | 0.020±0.0015 | 7.1±4.3 | 355 |
| NCH-51 | 3 | 9.4±0.15 | 149±28 | 16 |
| OKI-005 | 3 | 0.11±0.015 | 7.5±0.10 | 68 |

Results presented are mean GI<sub>50</sub> values +/- SEM. Relative resistance (RR) value was determined by dividing the GI<sub>50</sub> value for the MCF-7 DpVp300 cells by the GI<sub>50</sub> value for the MCF-7 cells.

**Supplemental Table S3.**
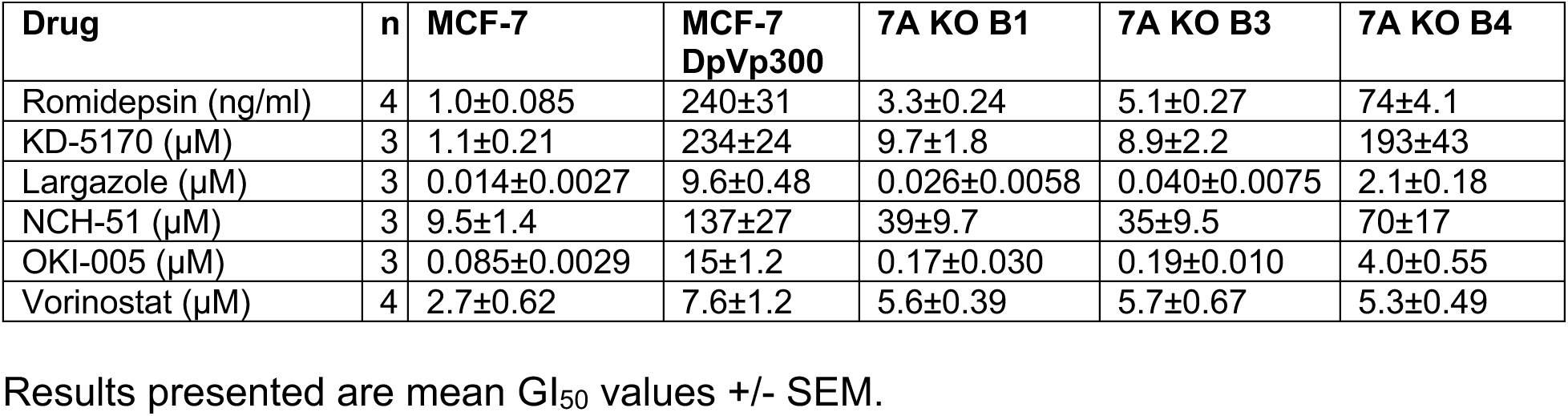
The cross resistance profile of MCF-7, MCF-7 DpVp300, 7A KO B1, 7A KO B3 and 7A KO B4 cells

| Drug | n | MCF-7 | MCF-7 DpVp300 | 7A KO B1 | 7A KO B3 | 7A KO B4 |
| --- | --- | --- | --- | --- | --- | --- |
| Romidepsin (ng/ml) | 4 | 1.0±0.085 | 240±31 | 3.3±0.24 | 5.1±0.27 | 74±4.1 |
| KD-5170 (µM) | 3 | 1.1±0.21 | 234±24 | 9.7±1.8 | 8.9±2.2 | 193±43 |
| Largazole (µM) | 3 | 0.014±0.0027 | 9.6±0.48 | 0.026±0.0058 | 0.040±0.0075 | 2.1±0.18 |
| NCH-51 (µM) | 3 | 9.5±1.4 | 137±27 | 39±9.7 | 35±9.5 | 70±17 |
| OKI-005 (µM) | 3 | 0.085±0.0029 | 15±1.2 | 0.17±0.030 | 0.19±0.010 | 4.0±0.55 |
| Vorinostat (µM) | 4 | 2.7±0.62 | 7.6±1.2 | 5.6±0.39 | 5.7±0.67 | 5.3±0.49 |

Results presented are mean GI<sub>50</sub> values +/- SEM.

**Supplemental Table S4.** The cross resistance profile of Vector, 7A 1-2, 7A 1-4, 7B 1-3 and 7B 2-2 cells

| Drug | n | Vector | 7A 1-2 | 7A 1-4 | 7B 1-3 | 7B 2-2 |
| --- | --- | --- | --- | --- | --- | --- |
| Romidepsin | 3 | 3.5±0.50 | 104±21 | 149±17 | 4.5±0.74 | 8.0±1.5 |
| KD-5170 | 5 | 4.2±1.0 | 134±12 | 133±9.7 | 40±7.1 | 47±13 |
| Largazole | 3 | 0.049±0.011 | 4.9±0.34 | 5.6±1.5 | 0.51±0.042 | 0.42±0.073 |
| NCH-51 | 3 | 7.5±1.7 | 103±12 | 71±17 | 182±47 | 117±31 |
| OKI-005 | 3 | 0.14±0.032 | 5.6±0.30 | 6.5±1.4 | 2.8±0.40 | 2.3±0.40 |
| Mocetinostat | 4 | 0.89±0.084 | 0.72±0.11 | 0.90±0.077 | 1.0±0.17 | 0.78±0.066 |
| Vorinostat | 3 | 1.1±0.19 | 0.90±0.076 | 1.4±0.12 | 1.2±0.088 | 1.2±0.058 |

Results presented are mean GI<sub>50</sub> values +/- SEM.

**Supplemental Table S5.** Normal tissue expression of METTL7A and METTL7B

| Tissue | METTL7A positive samples | METTL7B positive samples |
| --- | --- | --- |
| Brain | 9/14 | 9/14 |
| Testis | 15/15 | 15/15 |
| Prostate | 0/5 | 0/5 |
| Ovary | 0/7 | 0/7 |
| Breast | 5/5 | 0/5 |
| Cervix | 0/5 | 0/5 |
| Heart | 10/10 | 10/10 |
| Kidney | 15/15 | 15/15 |
| Liver | 13/14 | 13/14 |
| Lung | 0/15 | 0/15 |
| Pancreas | 0/5 | 0/5 |
| Esophagus | 0/5 | 0/5 |
| Stomach | 15/15 | 15/15 |
| Small intestine | 0/4 | 4/4 |
| Colon | 0/13 | 10/13 |
| Rectum | 0/5 | 1/5 |
| Skin | 3/3 | 3/6 |
| Lymph node | 0/5 | 0/5 |
| Spleen | 0/16 | 0/16 |
| Bone marrow | 0/5 | 0/5 |
| Thymus | 0/5 | 0/5 |

**Supplemental Table S6:** Staining methods and antibodies for immunohistochemistry

| Antibody | Dilution | Antigen Retrieval | Secondary Antibody kit | Control Tissue | Manufacturer | Catalog Number |
| --- | --- | --- | --- | --- | --- | --- |
| METTL7A | 1:500 | Citrate 20' | Bond polymer refine kit (minus postprimary reagent) | Empty vector and *METTL7A* transfected HEK-293 cells | Origene | TA346478 (rabbit polyclonal) |
| METTL7B | 1:500 | Citrate 20' | Bond polymer refine kit (minus postprimary reagent) | Empty vector and *METTL7B* transfected HEK293 cells | Sigma | HPA830644 (rabbit polyclonal) |

